# Microbial plankton in southern Hudson Bay and James Bay in late summer

**DOI:** 10.64898/2026.11.01.763523

**Authors:** Zakhar V. Kazmiruk, R. Eric Collins

## Abstract

Hudson Bay and James Bay, on the southern margin of the Arctic, are warming and freshening as river discharge increases and surrounding permafrost thaws, yet their microbial plankton remain poorly described, and James Bay and the waters around the Belcher Islands have never been surveyed with environmental DNA. We conducted the first dedicated DNA-based survey of eukaryotic and prokaryotic plankton in southern Hudson Bay and James Bay, sequencing 18S rRNA (V9) and 16S rRNA (V4) gene amplicons from 72 samples collected at 51 stations in August 2022 and 2023. Hierarchical clustering resolved nine distinct communities; the primary division separated surface James Bay from Hudson Bay, with further structuring by year, distance to coast, depth, and riverine influence. Temperature, salinity, depth, and distance to coast together explained 31% of community variation (distance-based redundancy analysis), and threshold indicator taxa analysis placed an abrupt community-level change in offshore surface waters at 55.3°N and a salinity of ∼24.5. Across this boundary, primary producers shifted from large *Chaetoceros* diatoms in Hudson Bay to the picoplanktonic green alga *Picochlorum* and the cyanobacterium *Cyanobium* in James Bay, while large *Gyrodinium* dinoflagellates and *Calanus* copepods gave way to *Gymnodinium*, *Pseudocalanus*, and small cercozoans and ciliates. Heterotrophic bacteria in Hudson Bay were dominated by Proteobacteria, Bacteroidota, and Verrucomicrobiota, likely using diatom- and dinoflagellate-derived organic matter, whereas Actinomycetota and Planctomycetota dominated James Bay, likely using cyanobacterial and terrestrial dissolved organic matter. Prokaryotic diversity was highest in riverine, estuarine, and James Bay waters and declined northward and seaward. We also report the first strong presence of marine *Synechococcus* CC9902 in Hudson Bay and identify cosmopolitan bacteria that provide a baseline for future monitoring. These results suggest that continued freshening of the Hudson Bay Complex will favour smaller primary producers and a stronger microbial loop, and that ecosystem shifts may occur abruptly once surface waters cross a salinity threshold.

## 1. Introduction

The Arctic Ocean is warming at nearly four times the global average (Rantanen et al., 2022). Over recent decades, Arctic has experienced accelerated permafrost thaw, river discharge, and reduced sea-ice extent (AMAP, 2024), all of which were previously shown to alter Arctic microbial plankton composition (Comeau et al., 2011; Delpech et al., 2021; Jackson et al., 2024). Because marine microorganisms form the base of aquatic food webs and drive carbon and nutrient cycling, these changes will have cascading impacts on biogeochemical cycles and food webs in the Arctic ecosystem (Sipler et al., 2017; Kellogg et al., 2019; Di Pane et al., 2024). Warming is also driving poleward shifts in marine taxa, including harmful algae, increasing ecological and food-safety concerns in northern waters (Anderson et al., 2021; McKenzie et al., 2021; Schiffrine et al., 2024; Husson et al., 2024). Despite their central role in ecosystem function, Arctic marine microorganisms remain poorly characterized, limiting our ability to understand and predict ecosystem changes (Royo-Llonch et al., 2021).

Hudson Bay Complex (HBC), defined as Hudson and James Bay and adjacent waters (Ungava Bay, Hudson Strait and Foxe Basin), is especially sensitive to climate changes due to its location on the southern margin of the Arctic, permafrost thaw, and large riverine discharge (Figure 1). The HBC was reported to exhibit a sea surface temperature anomaly of +2.6 °C in 2015-2024 relative to the 1900-2000 average (Husson et al., 2024). On the southern shores of HBC lie the Hudson Bay Lowlands, a vast permafrost peatland and boreal wetland that contains more carbon than Canada’s “managed” boreal forest (∼30 Pg; Harris et al., 2022; Li et al., 2025), at least half of which is predicted to thaw by 2100 (Gough and Leung, 2002). Finally, the annual runoff yield for Hudson and James Bays (∼0.9 m) is almost three times higher than that of the Arctic Ocean (Granskog et al., 2011) and continues to increase, freshening the region (MacDonald et al., 2018; Champagne et al., 2021). From 2011 to 2019, the river water fraction in surface waters of the Hudson and James Bays increased at a rate of 0.0048 yr^-1^, and sea ice melt fraction declined at -0.003 yr^-1^ (Basu et al., 2026). Hence, the HBC appears to be an ideal region for studying the accelerating impacts of warming, permafrost thaw, and rive discharge on the Arctic Ocean.

**Figure 1.**
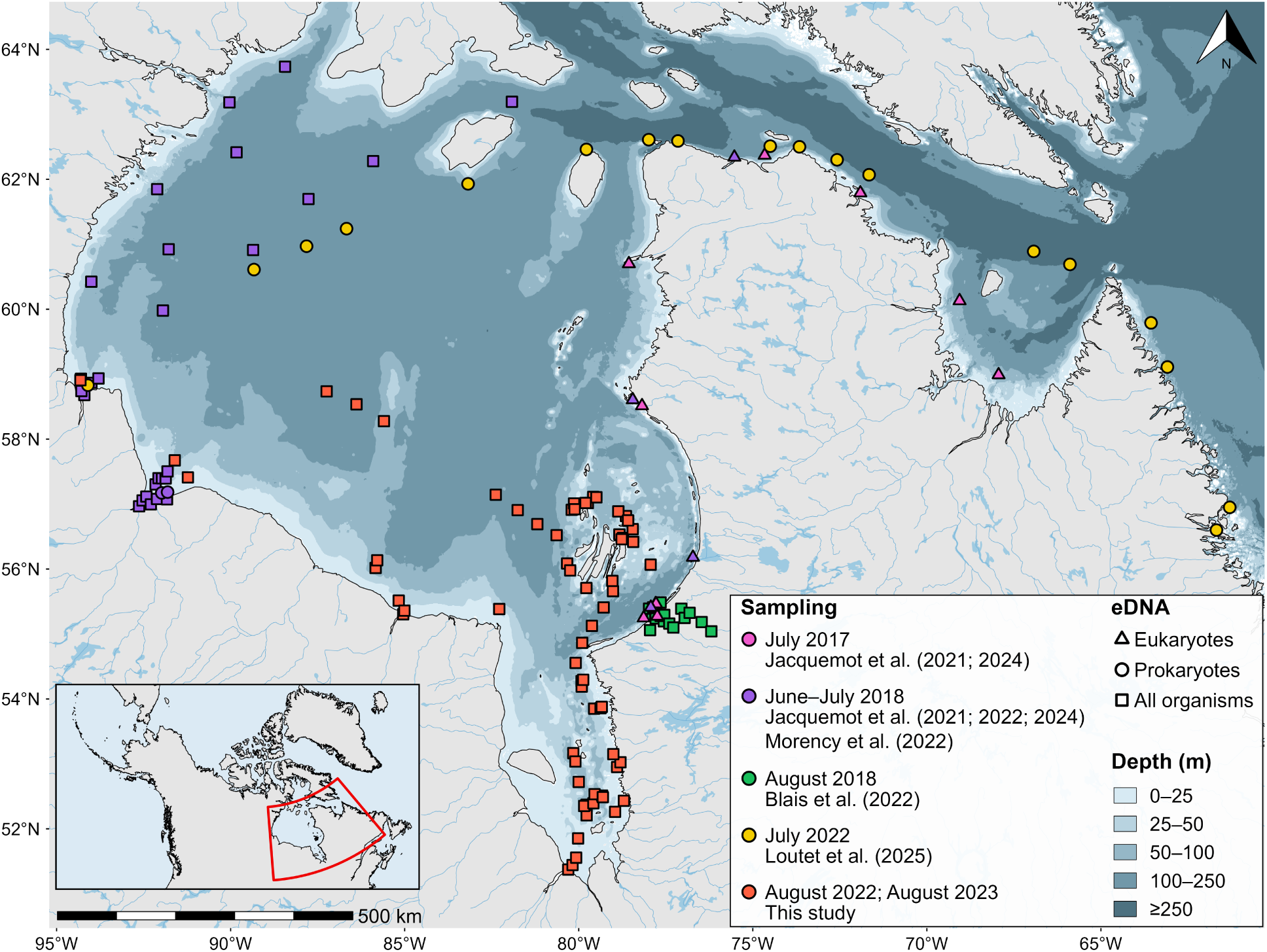
Environmental DNA (eDNA) sampling surveys in the Hudson Bay Complex during summer. Station colors indicate sampling periods and corresponding studies, while station symbols indicate the type of eDNA analyzed (eukaryotic, prokaryotic, or both). Foxe Basin is not shown due to absence of eDNA sampling. For nearby stations, 0.02° jitter was applied to prevent overplotting.

Within the HBC, James Bay deserves special attention due to a severe historical gap in the research of this region, despite its uniquely large riverine input, high vulnerability to permafrost thaw, strong downstream oceanographic influence, and concerns of the Indigenous communities residing in the area. James Bay is a large (67,000 km^2^) and shallow estuary-like body of water that receives 36% (323 km^3^ yr^-1^) of the total river discharge (McCullough et al., 2019). This massive riverine influx delivers high amounts of terrestrial dissolved organic matter (tDOM), particularly to the southeastern part of James Bay. Moreover, due to its southernmost location within the HBC, the region is particularly vulnerable to surrounding permafrost thaw, which further increases the tDOM input (Forget et al., 2025). Also, the large riverine input received by James Bay shapes the ecosystem around the Belcher Islands, located 120 km north of James Bay (Eastwood et al., 2020). In summer, James Bay water is transported to the Belcher Islands, where it gradually accumulates west of the islands and, due to cyclonic vorticity, also contributes to a persistent river water signal east of the islands (Basu et al., 2026). From there, the river water travels along eastern Hudson Bay and southern Hudson Strait into the North Atlantic Ocean (Straneo and Saucier, 2008). Finally, James Bay supports Cree First Nations, while the Belcher Islands host the Inuit community of Sanikiluaq. In recent decades, the local communities observed substantial environmental shifts, including altered river mouth morphology and changes in coastal flora and fauna (Dickey, 2015). Until recent field campaigns (Mundy et al., 2026), no dedicated offshore surveys had been conducted in James Bay since the 1970s (Peck, 1976).

Both freshwater and marine ecosystems depend on microscopic plankton, as they represent the base of the food chain and drive carbon and nutrient cycling. These organisms include primary producers such as cyanobacteria and algae, and heterotrophic bacteria that utilize dissolved and particulate organic matter. Heterotrophic protists and mesozooplankton serve as critical trophic links by feeding on microorganisms and particulate organic matter, channeling energy to higher trophic levels. Due to their abundance, diversity, and small size, oceanographers have started using DNA analysis to identify plankton and learn how they may be impacted by environmental changes. Recent environmental DNA (eDNA) surveys of Hudson Bay (Figure 1) show that summer microbial communities respond strongly to salinity and temperature. In western Hudson Bay, the change from ice-covered to open water conditions shifted phytoplankton communities from small green algae to large diatoms, which could increase the amount of food available to seafloor animals like clams (Jacquemot et al., 2022). In northern Hudson Bay, salinity and temperature strongly influenced the microbial community in surface waters (Loutet et al., 2025). In eastern Hudson Bay, where waters are stratified due to river runoff, mixotrophic dinoflagellates favoured the fresh surface mixed layer, while diatoms resided in colder subsurface waters (Jacquemot et al., 2024). In the river-estuary continuum of the Churchill, Nelson, and Great Whale Rivers, salinity mainly drove differences in microbial populations, while organic matter sources, hydrological variations, and estuarine circulation also contributed to shaping the microbial communities (Blais et al., 2022; Jacquemot et al., 2021; Morency et al., 2022). Overall, these studies suggest that future warming, freshening, and permafrost thaw in the Hudson and James Bay will fundamentally change microbial communities, with consequences for food webs and biogeochemical cycling. However, eDNA survey of microbial plankton around the Belcher Islands and in James Bay has never been done.

In this study, we conducted the first dedicated DNA-based survey of eukaryotic and prokaryotic microbial plankton in southern Hudson Bay and James Bay (SHJB; Figure 1). The objectives were twofold: (1) to describe later-summer plankton communities, and (2) to determine how environmental factors structure plankton composition across the region. Plankton compositions were derived via 16S-V4 and 18S-V9 rRNA gene amplicon sequencing. Potential environmental driver of the community changes were assessed. Based on prior Arctic and Hudson Bay studies, we hypothesized that late-summer plankton community composition in SHJB is driven by water-mass-defining physical parameters (temperature, salinity, and depth) and differs markedly between James Bay and Hudson Bay. This work is the microbiological component of a multidisciplinary effort to characterize the waters of James Bay and the Belcher Islands from 2021 to 2023 (Mundy et al., 2026).

## 2. Methods

### 2.1. Sampling and associated environmental data

Water samples were collected in southern Hudson Bay and James Bay in August 2022 (n = 38) and August 2023 (n = 34) as part of the Southern Hudson Bay and James Bay expeditions onboard the R/V *William Kennedy* (Figure 1). Between one and three depths were sampled at 51 stations (Stn), numbered chrono-logically. Each sample was characterised by latitude, longitude, depth, bottom depth, and sampling time. Temperature, salinity, chlorophyll-a fluorescence (fChl-a; measured by WET Labs ECO-AFL/FL fluorometer), CDOM fluorescence (fCDOM; measured by WET Labs CDOM fluorometer), and beam attenuation (WET Labs C-Star transmissometer, ∼650 nm) were measured concurrently and associated with each sample. Most samples were collected aboard the ship using 5-L Niskin bottles (n = 40), usually with a rosette equipped with a Sea-Bird CTD (n = 39). Because the rosette CTD lacked a transmissometer in both years and fluorometers in 2023, the missing optical measurements for most rosette samples (n = 36) were obtained from a standalone Sea-Bird CTD deployed on the same day, within 1.5 km of the sample location (median 0.4 km), and with a negligible difference in depth (median difference 0.13 m, range 0.02-0.24 m). Due to a rosette malfunction, the sample at Stn 44 (75 m) was taken with an individual Niskin bottle and paired with the measurements from a standalone Sea-Bird CTD taken on the same day, 0.4 km away, at a depth of 63 m. Additionally, samples were collected from the ship’s underway flowthrough system connected to a thermosalinograph (n = 27). For a more detailed environmental characterization, a subset of these samples (n = 11) was paired with the measurements from a standalone Sea-Bird CTD taken on the same day, within 1.5 km of the sample location (median 0.2 km), and at the same depth. The remaining flowthrough samples (n = 16) were paired with the thermosalinograph measurements corresponding in time and location. Finally, some samples were collected close to shore and within rivers from a Zodiac boat using a standalone Niskin bottle (n = 5). The sample at Stn 18 (1 m) was paired with the measurements taken by a standalone Sea-Bird CTD brought aboard the Zodiac, and the rest of the samples were paired with corresponding CastAway CTD measurements.

Two additional environmental parameters were derived. Mixed layer depth was calculated from downcast profiles for the 20 stations sampled by a rosette as the maximum depth at which potential density (sigma-theta) remained within 0.1 kg m^-3^ of the density measured at the shallowest depth (Peralta-Ferriz and Woodgate, 2015). Distance to coast and maps were generated using 1:10 m physical vectors from Natural Earth v5.1.1.

### 2.2. Sample processing and DNA sequencing

Within a few hours of sampling, the water collected was filtered using a peristaltic pump. Prior to the filtration, the tubing was rinsed with Milli-Q and sampled water. Filtration was performed through 0.22 µm Sterivex^TM^ filters until the filter clogged or showed visible color. Once filtration was done, filters were immediately stored at -20°C. At the University of Manitoba, the filters were thawed and immediately processed. Similar to the protocol by Cruaud et al. (2017), filter casings were opened with a sterilized PVC pipe cutter, and filters were transferred onto a sterile piece of foil, shredded using a sterile scalpel, and collected into PowerBead tubes. DNA was extracted using the DNeasy® PowerSoil® Pro Kit following the manufacturer’s protocol, quantified using a Qubit Flex Fluorometer, and stored at -80°C.

16S and 18S rRNA genes were amplified and sequenced with Illumina NovaSeq 6000 using the Earth Microbiome Project protocols (Caporaso et al., 2023; Amaral-Zettler et al., 2023). The V4 hypervariable region of the 16S rRNA genes in bacteria and archaea was amplified and sequenced using the 515F-806RB primer pair (Caporaso et al., 2011; Apprill et al., 2015). The V9 hypervariable region of the 18S rRNA gene in eukaryotes was amplified and sequenced using the 1391f-EukBr primer pair (Amaral-Zettler et al., 2009). The 250-bp paired end reads were processed using the R package “DADA2” v1.36 (Callahan et al., 2016) and the application “microscape” (https://github.com/rec3141/microscape), resulting in the generation of amplicon sequence variants (ASVs). The processing included demultiplexing, quality filtration, dereplication, and chimera removal.

### 2.3. Taxonomic Assignment

Prokaryotic taxonomic assignment was performed with the naive Bayesian classifier implemented with the R package “DADA2” v1.36 (Callahan et al., 2016) against the SILVA database (v138.2; Quast et al., 2013). At each taxonomic level, assignments with bootstrap support < 80% or those labeled “Unknown Family”, “Incertae Sedis”, or “Candidatus Nitrosoarchaeum” (https://www.arb-silva.de/documentation/release-1382/) were replaced with a placeholder, which indicates that the organism is unclassified (uncl.) at the level and includes the lowest resolved level (e.g., uncl. Dinophyceae). Sequences unclassified at domain level or classified as eukaryota, chloroplasts, and mitochondria were removed.

Eukaryotic taxonomic assignment was performed with the R package “rBLAST” v1.3.1 (Hahsler and Nagar, 2024) and the PR2 database (v5.1.1; Guillou et al., 2013). Only assignments with 100% query coverage (qcovhsp) and *≥* 80% identity (pident) were accepted. For each query sequence, the alignment with the highest bit score, highest identity, fewest mismatches, and fewest gap openings was used for taxonomic assignment. If multiple database sequences satisfied the selection criteria, we used the most detailed and frequent taxonomic assignment, preferably at the genus level. At each taxonomic level, assignments with prefix “_X” were removed and replaced with a placeholder denoting that the organism is unclassified (uncl.) at the level. Only query sequences identified as eukaryota were kept for further analysis, which effectively removed unmapped sequences and sequences mapped to organelles and prokaryotes. Taxonomic assignments of relevant taxa were manually verified using the core nucleotide database (core_nt) and RefSeq genome database (refseq_genomes) of the National Center for Biotechnology Information (Sayers et al., 2022). Overall, this workflow retained most query sequences suspected to be eukaryotic and ensured high-quality taxonomic assignments.

### 2.4. Statistical analyses

#### 2.4.1. Identification of distinct plankton communities

The original prokaryotic and eukaryotic ASV tables contained 159668 and 18014 ASVs, respectively. After removing rare ASVs (mean relative abundance across samples *≤* 0.01%), we retained 634 prokaryotic (620 bacteria, 14 archaea) and 434 eukaryotic ASVs, which together accounted for 97% and 89% of the total reads in each domain. These non-rare eukaryotic and prokaryotic ASVs were used to construct a combined ASV table, where relative abundances are expressed within each domain. All further analyses used this combined ASV table unless stated otherwise.

Dendrograms were constructed for both samples and ASVs by fourth-root-transforming the relative abundance table, calculating Bray-Curtis dissimilarity, and applying Ward’s D2 hierarchical clustering. The number of sample clusters was evaluated using silhouette widths and bootstrap resampling with Jaccard similarity over 1,000 iterations. Solutions ranging from 5 to 9 clusters were considered to capture the principal community patterns without over-splitting the data. The nine-cluster solution was the only evaluated solution in which every cluster had a bootstrap Jaccard similarity greater than 0.6 and was therefore selected for subsequent analyses. These clusters were considered stable under bootstrap resampling and represented samples with distinct microbial-community compositions. For visualization, sample branches were rotated toward a highest-to-lowest latitude order, while ASV branches were rotated according to abundance-weighted mean latitude. These rotations altered only the display order and did not change dendrogram topology, branch lengths, or cluster membership.

#### 2.4.2. Environmental characteristics and relationships with community composition

Environmental parameters were summarized by cluster using medians and ranges to account for skewed distributions (Table S1). Differences in environmental parameters among clusters were assessed with Kruskal-Wallis tests followed by post-hoc Dunn’s tests, where Benjamini-Hochberg (BH) adjusted p < 0.05 was considered significant (Table S2). Non-parametric tests were chosen because one-way ANOVA residuals violated normality assumptions (Shapiro-Wilk test, p < 0.05).

Influence of environmental parameters on microbial community composition was assessed using distance-based redundancy analysis (dbRDA) of Bray-Curtis dissimilarities calculated from the fourth-root-transformed combined ASV table, implemented with the function *dbrda* from the R package “vegan” v2.7.5 (Oksanen et al., 2026). Depth, distance to coast, temperature, and salinity were explanatory variables because they were available for all samples. The variables were standardized to zero mean and unit variance, and their low collinearity was confirmed via Pearson correlations and variance inflation factors below 5. Significance of the overall model, constrained axes, and marginal effect of each explanatory variable were verified using Monte Carlo tests (99,999 permutations). The variation explained by the model was reported as adjusted R².

Community-level change points along the principal geographic and hydrographic gradients separating Hudson Bay and James Bay were identified using Threshold Indicator Taxa Analysis (TITAN), implemented with the R package “TITAN2” v2.4.4 (Baker and King, 2010). Because riverine, coastal, and deep samples were unevenly represented across Hudson Bay and James Bay, TITAN was restricted to a surface pelagic subset (>10 km from the coast, 1-5 m depth; n = 39). Latitude and salinity were evaluated separately using default TITAN settings and retaining ASVs present in at least three samples.

#### 2.4.3. Alpha diversity across clusters and environmental gradients

Eukaryotic and prokaryotic Shannon diversity were calculated separately from the original 18S and 16S ASV count tables. Five samples with very low read counts were excluded, leaving 67 samples. As suggested by Schloss (2024), eukaryotic and prokaryotic libraries were independently rarefied 1,000 times to their respective minimum retained sequencing depths of 12,394 and 18,299 reads. Shannon diversity was calculated after each rarefaction, and the resulting values were averaged for each sample.

Differences in the Shannon diversity index between clusters were assessed with Kruskal-Wallis tests followed by post-hoc Dunn’s tests, where BH adjusted p < 0.05 was considered significant (Table S3). Non-parametric tests were chosen because one-way ANOVA residuals violated normality assumptions (Shapiro-Wilk test, p < 0.05). Associations between Shannon diversity and spatiotemporal and environmental parameters were assessed separately for eukaryotes and prokaryotes using Spearman rank correlations, where p < 0.001 was considered significant (Table S4).

#### 2.4.4. Indicator and cosmopolitan taxa

To identify taxa driving community shifts between clusters, we performed indicator taxa analysis in all samples (n = 72). Indicator ASVs (bioindicators) were identified using the function *multipatt* from the R package “indicspecies” v1.8.0 (De Cáceres and Legendre, 2009), which is based on the original method by Dufrêne and Legendre (1997). Individual clusters and pairs of clusters were considered as potential target groups. For each ASV, the specificity (A), fidelity (B), and permutation-based p-value were evaluated for the same best supported cluster or pair of clusters identified. When an ASV was significantly associated with a pair of clusters, it was assigned for visualization and interpretation to the individual cluster within that pair, in which it had the highest mean relative abundance. To retain representative bioindicators, criteria stricter than those of Dufrêne and Legendre (1997) were applied: at least 60% of the ASV’s summed group-specific mean relative abundance had to occur in the target group (A *≥* 0.6), the ASV had to occur in at least 60% of samples in the target group (B *≥* 0.6), and the association had to be significant (p < 0.01).

To identify ecologically flexible taxa, cosmopolitans were detected among the ASVs that were not bioindicators for any cluster or pair of clusters. An ASV was classified as a cosmopolitan when it occurred in at least 60% of samples in each of the eight non-riverine clusters.

To assess potential contamination in the flowthrough system, we applied the same indicator taxa criteria (A *≥* 0.6, B *≥* 0.6, and p < 0.01) to offshore surface samples (n = 39; distance to coast >10 km; depth 1-5 m) to identify ASVs specifically associated with the flowthrough system (n = 25) rather than the rosette (n = 14). No ASVs were significantly associated with flowthrough samples, providing no evidence of systematic contamination attributable to the flowthrough system.

### 2.5. Software and data availability

Analyses and visualizations were performed in R version 4.6.1. The complete analysis pipeline and list of packages used are available at https://github.com/kazmiruz/WK2022-23. The input files required to run the analysis pipeline and output files are available at https://osf.io/r32sy. Raw sequence reads and metadata are deposited in the NCBI Sequence Read Archive under BioProject PRJNA421293 (BioSamples SAMN62001260-SAMN62001331).

## 3. Results

### 3.1. Distinct plankton community clusters

By applying Bray-Curtis dissimilarity and Ward’s hierarchical clustering to the relative abundance of non-rare eukaryotic and prokaryotic ASVs, samples from southern Hudson Bay and James Bay were partitioned into nine clusters, each representing a distinct plankton community (Figure 2; Figure S1). The clusters were named according to their year (2022 and 2023), water layer (surface and subsurface), and geographic location (Figure 2). Planktonic communities in surface waters of southern Hudson Bay are represented by clusters oHB and BI22 sampled in 2022, and clusters cHB and EST sampled in 2023. oHB was sampled offshore, exclusively via the flowthrough system, while BI22 encompasses the waters around the Belcher Islands. cHB is located in coastal Hudson Bay, including the area around the Belcher Islands, while EST represents the Churchill and Winisk estuaries. Planktonic communities in surface waters of northern and southern James Bay are represented by cluster nJB, sampled in 2022, and cluster sJB, sampled in both 2022 and 2023, respectively. MR is the only riverine cluster and represents the Maquatua River located on the northeastern coast of James Bay. Finally, planktonic communities in the subsurface layer are represented by cluster dBI around the Belcher Islands, sampled in 2022 and 2023, and cluster dJB in James Bay, sampled in 2022.

**Figure 2.**
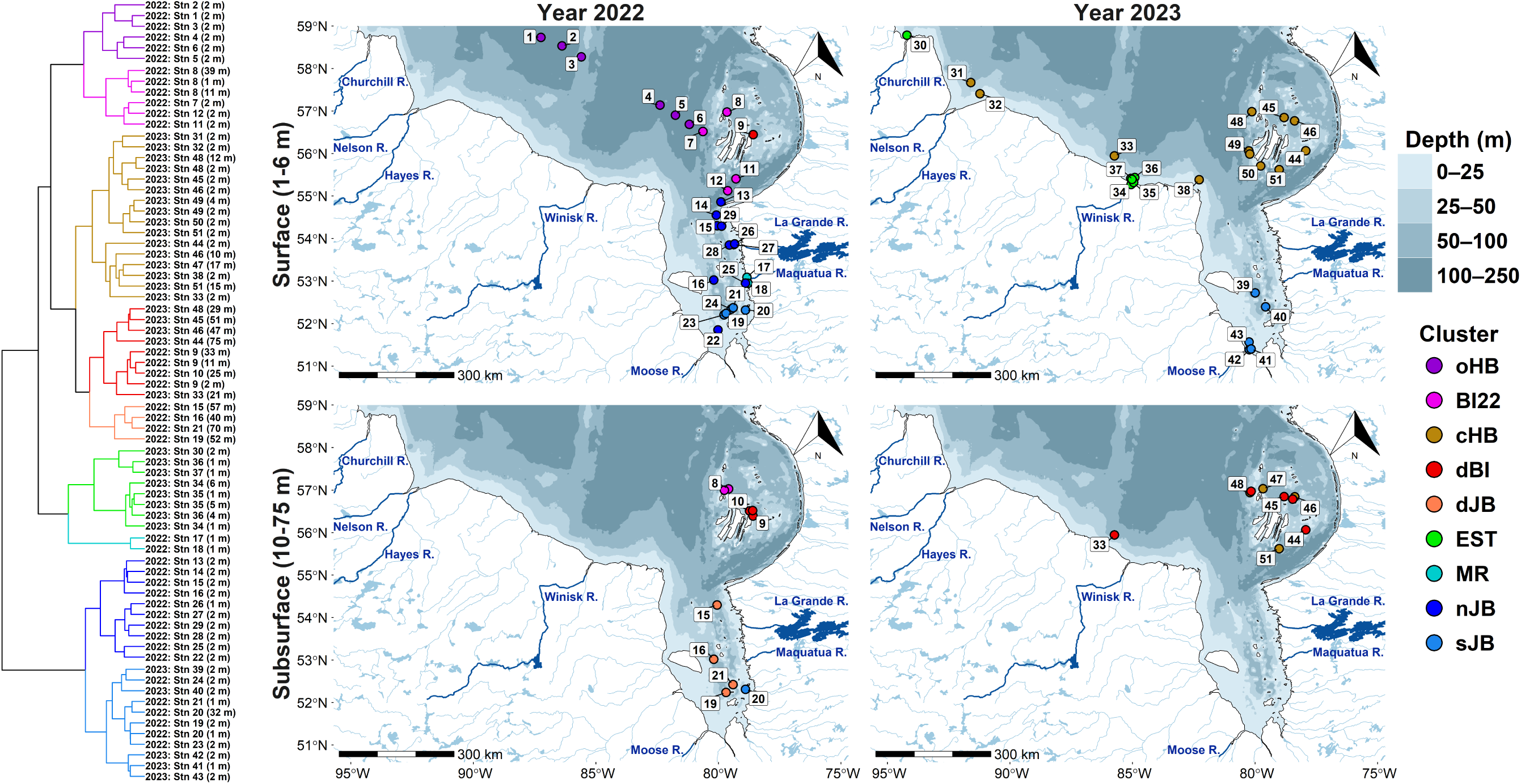
Environmental DNA samples collected in this study in southern Hudson Bay and James Bay. Maps show surface and subsurface samples collected in August 2022 and August 2023 as circles, colored by cluster. Boxed numbers indicate stations. For nearby samples, 0.1° jitter was applied to prevent overplotting. The corresponding cluster dendrogram is shown on the far left: branches are colored by cluster, and sample names indicate year, station (Stn), and depth.

The hierarchical clustering reveals a clear segregation in plankton between Hudson Bay and James Bay, with additional segregation by year, distance to coast, and depth (Figure 2; Figure S1). The main split separates the surface James Bay (nJB and sJB) from all remaining clusters. The next major split isolates the Churchill and Winisk River estuaries (EST) and the Maquatua River (MR) from surface Hudson Bay and all the deep waters. The remaining six clusters then split into three groups: the offshore and Belcher Islands waters sampled in 2022 (oHB and BI22), the coastal Hudson Bay waters sampled in 2023 (cHB; includes waters around the Belcher Islands), and deep waters of both Hudson Bay and James Bay (dBI and dJB).

### 3.2. Environmental characteristics of clusters

Spatial and water-mass-defining parameters (depth, distance to coast, temperature, and salinity) were available for all clusters and differed significantly among them (Tables S1 and S2; Figure 3A). dBI was significantly deeper (median depth 29.0 m) than all clusters except BI22 and dJB. The lack of a significant difference between dBI and BI22 likely reflects the relatively deep mixed layers at Stn 8 (14.4 m) and Stn 9 (17.8 m), where all samples – even those collected below the mixed layer – were assigned to BI22 and dBI, respectively. dJB was significantly deeper (median depth 54.3 m) than all other clusters except dBI. oHB was significantly farther from the coast (median distance 191 km) than all clusters except dJB. Among surface clusters, EST covered the widest salinity range (15.3-27.5), while MR, nJB, and sJB were significantly less saline (*≤* 24.2) than oHB, cHB, and BI22. dJB was significantly more saline than one of the corresponding surface clusters, sJB. BI22 was significantly colder (median temperature 4.1 °C) than all other surface clusters except oHB. Subsurface clusters, dBI and dJB, were significantly colder (median temperature 2.9 °C) than cHB, EST, MR, nJB, and sJB. No significant differences were detected between nJB and sJB or between dBI and dJB.

**Figure 3.**
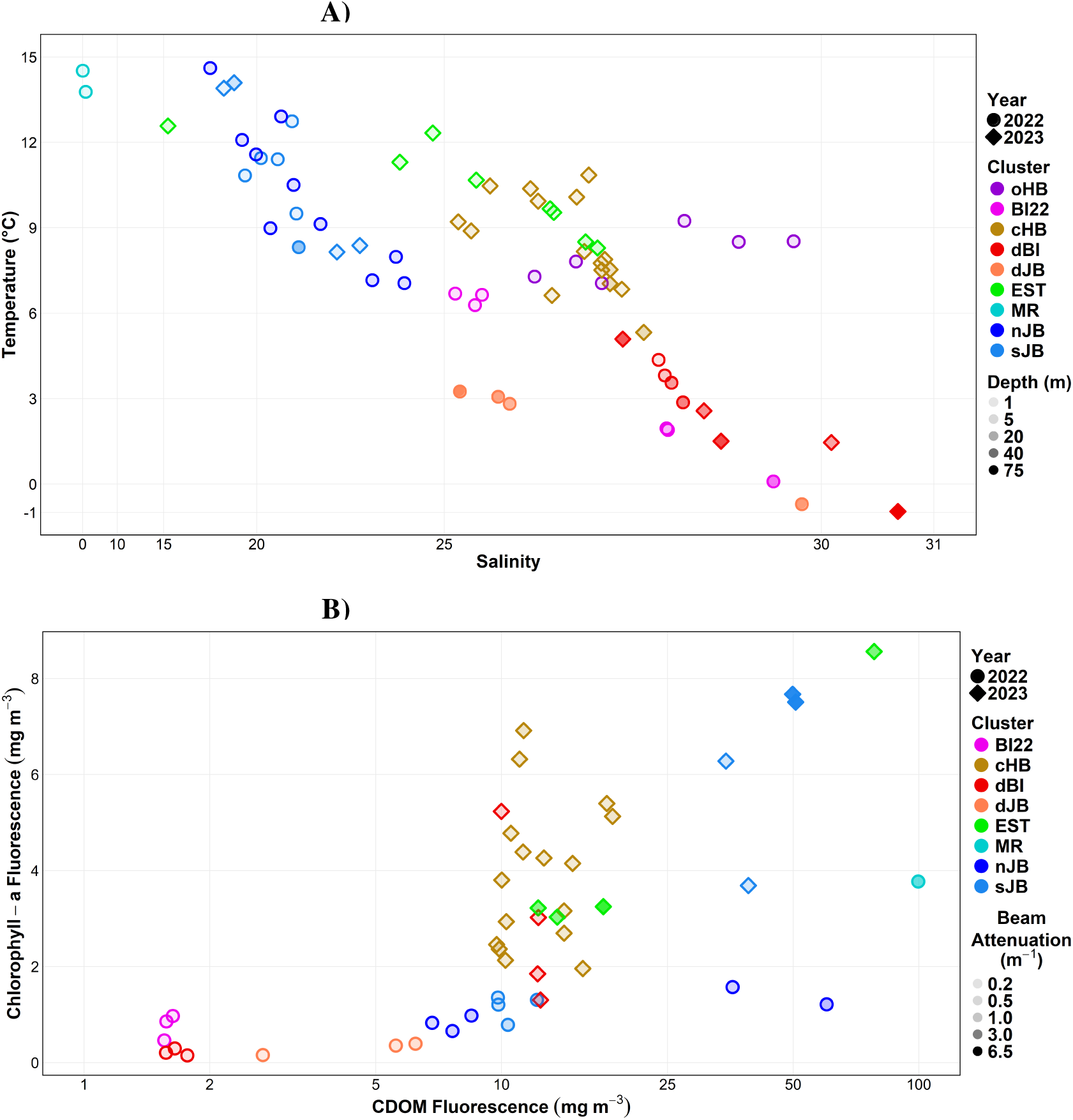
Environmental parameters across clusters. A) Relationship between temperature, salinity, and depth. B) Relationship between chlorophyll-a fluorescence, CDOM fluorescence, and beam attenuation. Sample points are shaped by sampling year and colored by cluster.

Optical parameters (fChl-a, fCDOM, and beam attenuation) were available for all clusters except oHB and differed significantly among clusters (Tables S1 and S2; Figure 3B). In BI22, dBI, dJB, EST, MR, nJB, and sJB, at least one optical parameter was missing for some samples, although measurements were available for at least half of the samples in each cluster. Values of fChl-a were significantly higher in cHB than in dBI and dJB, and in EST and sJB than in dJB. The highest values were in surface waters (*≤* 2 m) near the Churchill River (8.6 mg m^-3^ at Stn 30) and the Moose River (7.5-7.7 mg m^-3^ at Stns 41-43). Values of fChl-a were also elevated in surface and intermediate waters (*≤* 17 m) around the Belcher Islands in 2023 (3.8-6.9 mg m^-3^ at Stns 44-51). Values of fCDOM were significantly higher in EST and sJB than in BI22, dBI, and dJB; in MR than in BI22 and dJB; and in cHB than in BI22. The highest values were in the Maquatua River (100 mg m^-3^ at Stn 18), Churchill River estuary (78 mg m^-3^ at Stn 30), and in James Bay near the Maquatua and Moose Rivers (50-60 mg m^-3^ at Stns 25 and 41-43). Beam attenuation was significantly higher in EST than in BI22, cHB, dBI, and nJB; in sJB than in BI22, cHB, and dBI; and in MR than in BI22. The highest values were near the Moose River (5.7-6.1 m^-1^ at Stns 41-43). Beam attenuation was also elevated in the Maquatua River and in the Churchill and Winisk River estuaries (2.6-3.8 m^-1^ at Stns 18, 30, 34, and 35).

### 3.3. Environmental relationships with plankton community composition

dbRDA was performed to resolve the influence of temperature, salinity, depth, and distance to coast on plankton community structure in the clusters (Figure 4). Overall, the model explained 31% of the total variance (adjusted R^2^ = 0.31). Temperature and salinity were anticorrelated along the primary axis, dbRDA1, which explained 18.0% of the variation. Distance to coast was positioned mainly along dbRDA2, which explained 8.2% of the variance. Depth and temperature were anticorrelated along dbRDA3, which explained 6.0% of the variance. Distance to coast was nearly orthogonal to the other three parameters.

**Figure 4.**
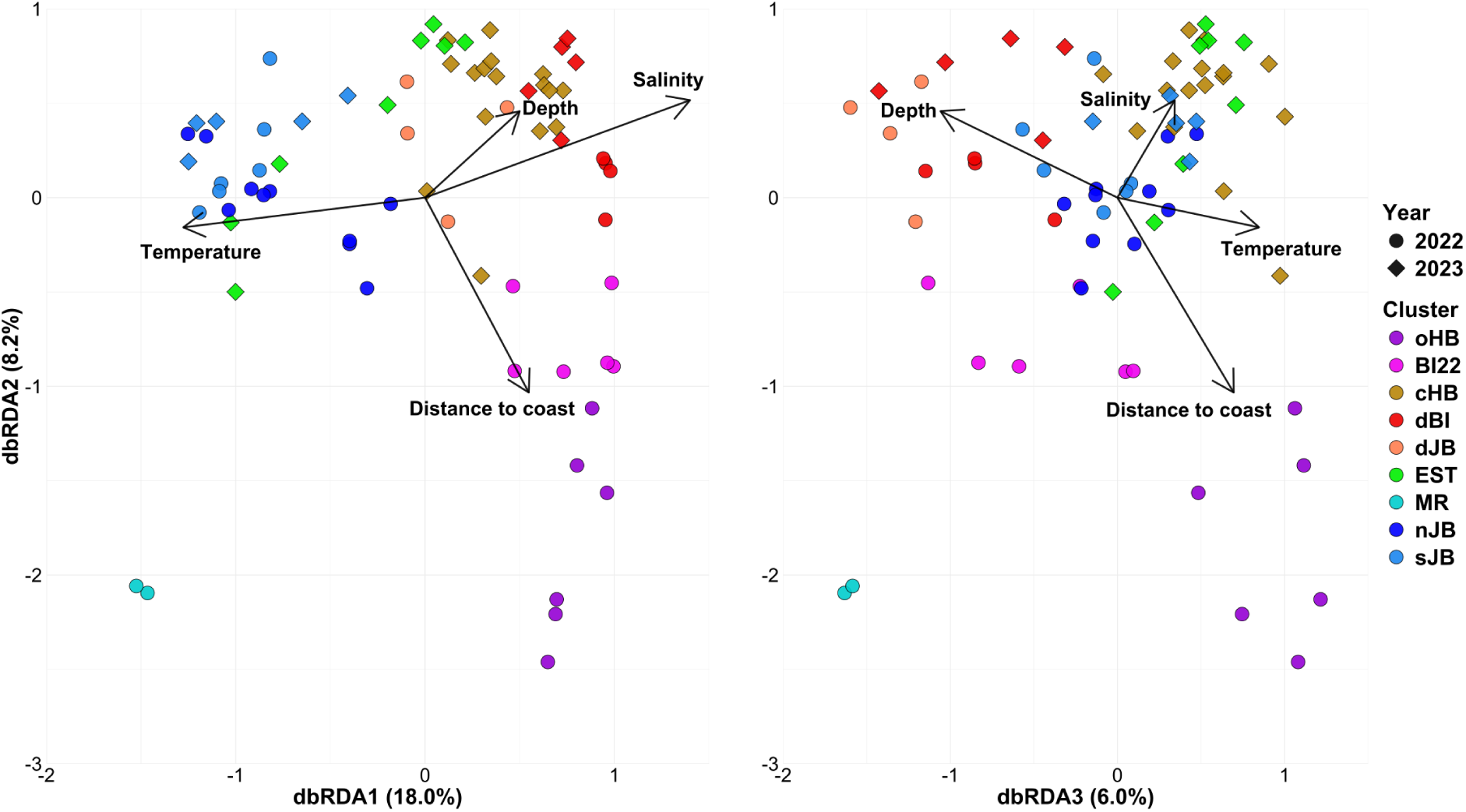
dbRDA ordination plot of plankton communities. Sample points are shaped by sampling year and colored by cluster. Vectors represent explanatory variables. had no significant correlation (Table S4). These trends demonstrate that prokaryotic diversity declined northward and seaward in SHJB.

MR was separated from the marine clusters and associated with low salinity. nJB and sJB were separated from the rest of the marine clusters and associated with low salinity and high temperature. oHB and BI22 were positively associated with distance to coast, with BI22 additionally associated with low temperature. cHB was associated with high salinity. EST was consistently associated with close proximity to coast and shallow depth and spanned the transition between coastal Hudson Bay and James Bay, partially clustering with both cHB and nJB/sJB. Finally, both dBI and dJB were associated with deep, cold waters, but dJB exhibited an association with greater depth and lower salinity than dBI.

TITAN was performed to identify the change points in latitude and salinity associated with abrupt shifts in offshore surface plankton composition. Offshore surface samples ranged from 51.5°N to 58.7°N and from salinities of 17.9 to 29.7. Community-level change points occurred at 55.3°N and around salinity 24.5.

### 3.4. Plankton composition and alpha diversity across clusters

#### 3.4.1. Alpha diversity across clusters and environmental gradients

Eukaryotic and prokaryotic diversity dynamics in southern Hudson Bay and James Bay were assessed using the Shannon diversity index. In this study, targeting the shorter eukaryotic V9 region (∼130 bp) versus the prokaryotic V4 region (∼250 bp) likely contributes to lower taxonomic resolution and fewer detected ASVs in eukaryotes. Additionally, the DADA2 pipeline removes singletons, decreasing observed richness. However, the impact on the Shannon diversity index is minimal as the metric accounts for both richness and evenness. First, eukaryotic diversity varied statistically significantly between clusters, where southern James Bay exhibited the lowest Shannon diversity index (Figure 5). Specifically, sJB had significantly lower diversity than cHB, dBI, and EST (Table S3). Spearman’s test did not reveal a statistically significant correlation between eukaryotic diversity and any spatiotemporal and environmental parameters. Second, prokaryotic diversity varied significantly between clusters, where the riverine, estuarine, and James Bay samples generally had a high Shannon diversity index (Figure 5). Specifically, dJB, EST, MR, and sJB had significantly higher diversity than oHB, BI22, and cHB (Table S3). Moreover, prokaryotic diversity correlated negatively with distance to the coast, latitude, and bottom depth, while other spatiotemporal and environmental parameters

**Figure 5.**
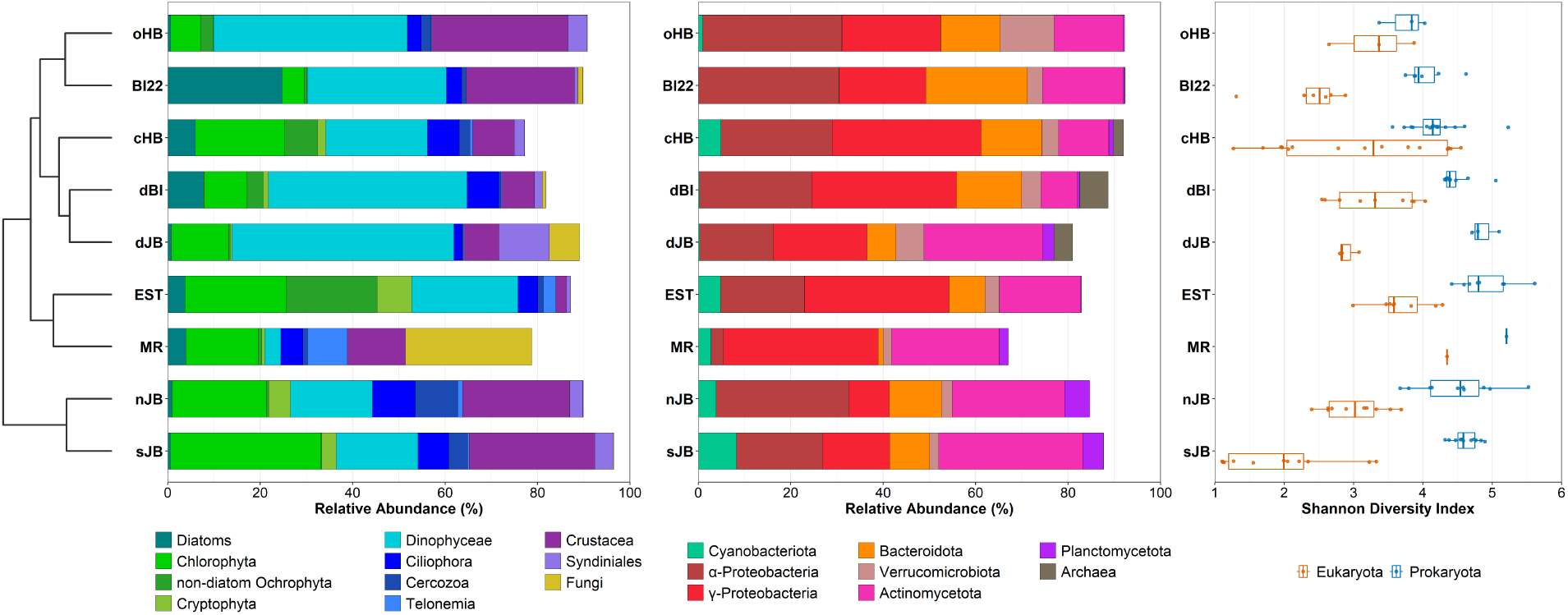
Overview of plankton communities in SHJB. For each cluster, major taxonomic groups and Shannon diversity are shown for both eukaryotes and prokaryotes. The cluster dendrogram is displayed on the far left.

#### 3.4.2. Photosynthetic algae and non-parasitic dinoflagellates

Photosynthetic algae were mainly represented by diatoms, Chlorophyta, non-diatom Ochrophyta, Cryptophyta, and Haptophyta (Figure 5). Diatoms (Mediophyceae, Coscinodiscophyceae, and Bacillariophyceae) were relatively abundant around the Belcher Islands, with the three most abundant genera being *Chaetoceros*, *Minidiscus*, and *Arcocellulus* (Figure S2A). Diatoms peaked in BI22 (25%), where *Chaetoceros* (ASV_918) was the sole phototrophic bioindicator. Diatoms were rarer in dBI (6%), cHB (8%), MR (4%), and EST (4%). Other bioindicator diatoms were *Minidiscus* (ASV_752) and *Arcocellulus* (ASV_946) for cHB, and *Aulacoseira* (ASV_925) for MR. Non-diatom photosynthetic algae were mainly present in river-influenced waters, with the three most abundant genera being *Picochlorum*, *Micromonas* and *Aureococcus* (Figure S2B). Chlorophyta (green algae) dominated this group, accounting for 33% of the relative abundance in sJB and 20-24% in cHB, EST, and nJB. Among bioindicator green algae, *Micromonas* favoured dBI (ASV_770) and EST (ASV_695); *Picochlorum* (ASV_680) preferred sJB, reaching 51% near the Moose River; and *Choricystis* (ASV_714) was restricted to MR. Non-diatom Ochrophyta and Cryptophyta peaked in EST (20% and 7%), with *Aureococcus* (ASV_641) and *Plagioselmis* (ASV_725) as bioindicators. Haptophyta were rare (∼2% in dBI, cHB, and EST), with *Phaeocystis* (ASV_761) as a bioindicator of dBI.

Dinophyceae (non-parasitic dinoflagellates) prevailed in offshore and deep waters (42-48% in oHB, dBI, and dJB) and were relatively abundant in other non-riverine clusters (18-30%) (Figure 5). The three most abundant genera of Dinophyceae were *Gyrodinium*, *Heterocapsa*, and *Gymnodinium* (Figure S2C). Among bioindicator Dinophyceae, *Gyrodinium* (ASV_945) and *Karlodinium* (ASV_818) represented oHB, *Biecheleria* (ASV_798) represented cHB, *Gyrodinium* (ASV_635) represented dBI, *Heterocapsa* (ASV_1061) and *Warnowia* (ASV_1026) represented EST, *Prorocentrum* (ASV_1034) represented dJB, and *Gymnodinium* (ASV_759) represented sJB. According to the NCBI BLAST Tree View, ASV_1061 belongs to a monophyletic branch of *Heterocapsa rotundata*.

#### 3.4.3. Heterotrophic eukaryotes

Heterotrophic eukaryotes were heterotrophic protists, zooplankton, parasitic dinoflagellates, and fungi (Figure 5). Among heterotrophic protists, Ciliophora, Cercozoa, and Telonemia were most abundant in riverinfluenced waters. Ciliophora reached 9% in nJB and 7% in cHB, dBI, and sJB; Cercozoa peaked in nJB (9%) and sJB (4%); Telonemia increased sharply in MR (8%). Key bioindicator heterotrophic protists included *Cyrtostrombidium* (ASV_815) for BI22, *Calliacantha* (ASV_974) for cHB, *Mesodinium* (ASV_726) for MR, *Ebria* (ASV_661) for nJB, and *Mesodinium* (ASV_668) for sJB. Zooplankton were mainly Crustacea, which were most abundant in oHB, BI22, nJB, and sJB (23-29%). *Calanus* (ASV_957) was a key bioindicator for BI22, and *Pseudocalanus* (ASV_889) for nJB. Parasitic dinoflagellates Syndiniales were mainly represented by Dino-Group-I (DG-I) and Dino-Group-II (DG-II), with DG-I-Clade-4 (ASV_954) as a bioindicator for oHB, DG-I-Clade-1 (ASV_979) and DG-II-Clade-10-and-11 (ASV_638) as bioindicators of dJB, and DG-I-Clade-4 (ASV_659) as a bioindicator for sJB. Fungi were most strongly present in MR (27%) and dJB (7%), where *Penicillium* (ASV_1021) was a bioindicator for MR, and *Rhodotorula* (ASV_734) and *Graphium* (ASV_693) for dJB.

#### 3.4.4. Cyanobacteria

Cyanobacteria were relatively abundant in the surface waters of SHJB, representing 8% of the prokaryotic community’s abundance in sJB, 4-5% in other river-influenced waters (cHB, EST, and nJB), and 3% in MR (Figure 5). Among the 22 non-rare cyanobacterial ASVs detected, 21 belonged to the order Synechococcales, which were classified as either *Synechococcus* CC9902 (2 ASVs) or *Cyanobium* PCC-6307 (19 ASVs). Furthermore, bioindicator analysis revealed cyanobacterial partitioning along the salinity gradient and between the two river-influenced regions (Figures 6 and S2D). *Synechococcus* CC9902 (ASV_17) characterised cHB, where it accounted for 99% of Synechococcus relative abundance and exceeded 10% prokaryotic relative abundance at salinities of 27.3-27.6. In contrast, *Cyanobium* PCC-6307 (ASV_210) characterised sJB, where it accounted for 55% of *Cyanobium* relative abundance and exceeded 10% prokaryotic relative abundance at salinities of 17.9-21.2. Lastly, *Cyanobium* PCC-6307 (ASV_221) characterised EST, where it accounted for 42% of *Cyanobium* relative abundance. ASV_221 also appeared in MR and some offshore waters.

**Figure 6.**
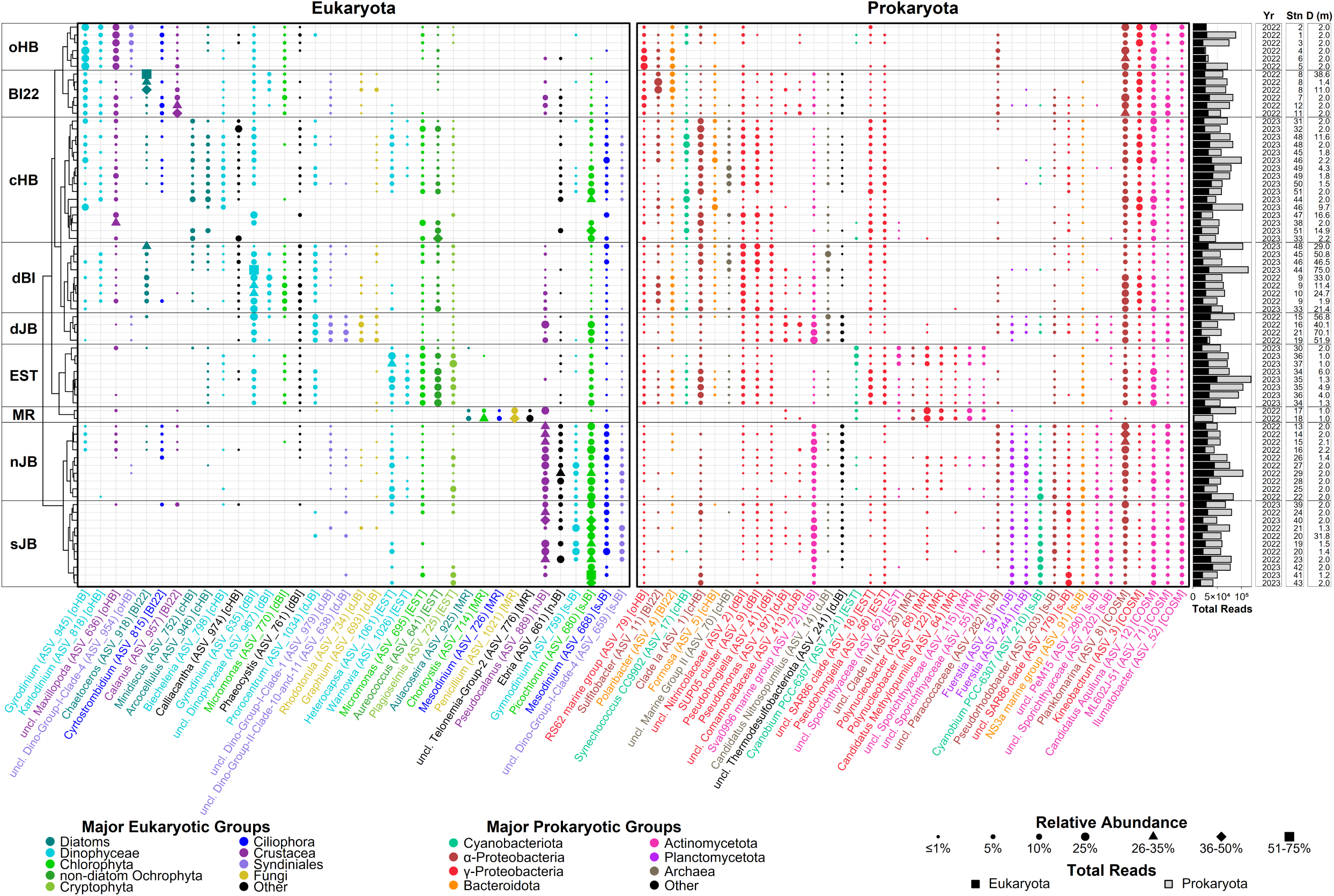
Eukaryotic and prokaryotic bioindicators and cosmopolitans in SHJB. Only ASVs with *≥*1.5% mean relative abundance within their indicator cluster (or across all samples for cosmopolitans) are shown. Colors indicate major taxonomic groups; square brackets show cluster affinity (COSM = cosmopolitan). For each sample, cluster and dendrogram order are shown on the left, and sequencing depth, year, station, and depth are provided on the right.

Given the polyphyletic nature of Synechococcales (Strunecký et al., 2023), we carefully verified the identities of the mentioned cyanobacterial ASVs. The best BLASTn alignments of *Synechococcus* CC9902 (ASV_17) against the NCBI RefSeq genome database of cyanobacteria (100% coverage and 98.4-99.6% identity) were exclusively marine species. The best BLASTn alignments (100% coverage and 98.4-100% identity) against the NCBI RefSeq genome database supported divergent ecological niches of the two *Cyanobium* bioindicators: the James Bay-specific ASV_210 aligned primarily with estuarine species, whereas estuarine and riverine ASV_221 aligned exclusively with freshwater species.

#### 3.4.5. Heterotrophic and chemotrophic prokaryotes

Heterotrophic and chemotrophic prokaryotes were mainly represented by *α*-and *γ*-Proteobacteria, Bacteroidota, Verrucomicrobiota, Actinomycetota, Planctomycetota, and Archaea (Figure 5). *α*-and *γ*-Proteobacteria were dominant and together accounted for 33-56% of the prokaryotic relative abundance across all clusters. Specifically, *α*-Proteobacteria comprised ∼30% in oHB, BI22, and nJB, while *γ*-Proteobacteria accounted for ∼30% in cHB, EST, MR, and dBI. Based on the bioindicator analysis, in Hudson Bay, oHB was characterised by *RS62 marine group* (ASV_79); BI22 by *Sulfitobacter* (ASV_11); cHB by Pelagibacterales Clade Ia (ASV_1); and dBI by SUP05 cluster (ASV_9) and *Pseudohongiella* (ASV_41). In James Bay, several brackish and freshwater taxa were bioindicators: MR was characterised by *Pelagibacterales* Clade III (ASV_29), *Polynucleobacter* (ASV_68 and ASV_122) and *Candidatus* Methylopumilus (ASV_64); sJB by *Pseudorhodobacter* (ASV_203); and dJB by *Pseudomonas* (ASV_197). Finally, *Planktomarina* (ASV_8) and *Kineobactrum* (ASV_3) were cosmopolitan.

Among other bacteria, Bacteroidota and Verrucomicrobiota were abundant relative to other prokaryotes in Hudson Bay waters, while Actinomycetota and Planctomycetota were more relatively abundant in James Bay waters (Figure 5). Bacteroidota and Verrucomicrobiota peaked at 22% in BI22 and 12% in oHB, respectively. Bacteroidota were mainly Flavobacteriales, with bioindicators *Polaribacter* (ASV_4) for BI22, *Formosa* (ASV_5) for cHB, and NS3a marine group (ASV_91) for sJB. While no prevalent bioindicators were identified for Verrucomicrobiota, this phylum was most abundant in oHB, where *Roseibacillus* accounted for 76% of the phylum’s relative abundance. In James Bay clusters (dJB, MR, nJB, sJB), Actinomycetota and Planctomycetota constituted 23-31% and 2-5%, respectively. Key bioindicator Actinomycetota were Sva0996 marine group (ASV_72) for dJB; multiple Sporichthyaceae taxa for EST, MR, and sJB; and PeM15 (ASV_290) for sJB. Among Planctomycetota, *Fuerstia* (ASV_154 and ASV_244) were identified as bioindicators for nJB. Finally, *Candidatus* Aquiluna (ASV_12) and *Ilumatobacter* (ASV_52) were cosmopolitan.

Lastly, Archaea peaked in relative abundance in deep waters, 6% in dBI and 4% in dJB. *Candidatus* Nitrosopumilus (ASV_14) was a key bioindicator for dJB.

## 4. Discussion

### 4.1. Surface pelagic waters

Summer plankton communities in the surface pelagic waters of James Bay were distinct from those in Hudson Bay. Hierarchical clustering based on ASV relative abundance clearly separated the James Bay surface clusters (sJB and nJB) from the remaining clusters (Figure 2). The dbRDA supported this pattern, showing that low salinity and high temperature separated the James Bay surface clusters from the remaining clusters along the primary axis (Figure 4). Finally, within the surface pelagic subset, TITAN identified community-level change points at 55.3°N and near salinity 24.5, coinciding with the geographic and hydrographic separation between James Bay and Hudson Bay (Figures 1b and 2a). The salinity-dependent segregation of summer surface plankton communities appears to be a broader feature of the HBC. Jacquemot et al. (2024) similarly found that salinity and temperature separated coastal summer protist communities of eastern Hudson Bay from those of the combined Hudson Strait and Ungava Bay region. Loutet et al. (2025) also found that salinity and temperature significantly affected summer prokaryotic community structure in surface waters of northern Hudson Bay and Hudson Strait.

On a smaller scale, surface pelagic waters of both Hudson Bay and James Bay were represented by a mosaic of distinct plankton communities that were shaped by the local environment (Figure 2). According to dbRDA, plankton communities in surface water of Hudson Bay were strongly influenced by distance to coast and temperature change around the Belcher Islands between 2022 and 2023 (Figure 4). oHB was a truly offshore Hudson Bay community (Table S1), where the orthogonal position of distance to coast to the other physical parameters suggested that the plankton community in the cluster was governed by non-physical factors. BI22 consisted of the cold waters sampled around the Belcher Islands in 2022, while cHB encompassed all coastal waters in 2023, both along the continental coast and around the Belcher Islands (Figure 3A; Table S1). Likewise, surface waters of James Bay comprised two pelagic clusters – northern (nJB) and southern (sJB) – with the division occurring near Akimiski Island. Both clusters were characterised by low salinity and high temperature (Figure 3A; Table S1), and the dbRDA did not separate them along the measured physical gradients (Figure 4), implying that non-physical factors drove the community differences within James Bay.

#### 4.1.1. Phytoplankton

In the surface pelagic waters of SHJB, photosynthetic algae and non-parasitic dinoflagellates differed substantially between Hudson Bay and James Bay, reflecting a transition from marine to river-influenced conditions, respectively. Hudson Bay (oHB, BI22, and cHB) was characterised primarily by diatoms and large dinoflagellates, while James Bay (sJB and nJB) was preferred by green algae and, to a lesser extent, small dinoflagellates (Figure 5). First, diatoms were relatively abundant around the Belcher Islands in both 2022 and 2023 (Figure S2A). *Chaetoceros* favoured the cold well-mixed waters of BI22 north of the islands, while *Minidiscus* and *Arcocellulus* characterised cHB, reaching higher abundance in warm waters situated along the James Bay riverine outflow east of the islands (Figure 6). While reported to co-dominate in high latitude waters such as the Russian Arctic Seas (Belevich et al., 2019) and Northwest Atlantic (Robicheau et al., 2022), these diatoms vary in size from chain-forming *Chaetoceros* (5-30 µm) to smaller *Arcocellulus* (∼5 µm) and *Minidiscus* (<5 µm) (Kaczmarska et al., 2009; Ribeiro et al., 2020). Moreover, unlike *Minidiscus* and *Arcocellulus*, *Chaetoceros* increased in relative abundance with depth, from 34% at 1.4 m to 73% at 38.6 m at Stn 8. This relative accumulation of *Chaetoceros* below the 14.4 m mixed layer depth suggests an efficient vertical carbon export mediated by these large chain-forming diatoms. Second, green algae and other non-diatom photosynthetic algae reached their highest relative abundance in James Bay (Figure 6 and S2B). They were also strongly represented in coastal Hudson Bay and south of the Belcher Islands, likely due to freshwater inputs from the southern Hudson Bay rivers and James Bay outflow. sJB was characterised by *Picochlorum*, which reached a relative abundance of over 50% near the Moose River. Interestingly, *Picochlorum* was not previously reported in the HBC, although this genus was recently documented in the river-influenced area south of the coast of Newfoundland in August, where it was more relatively abundant than other green algae (Mordret et al., 2024). Unlike green algae, Cryptophyta were more relatively abundant in nJB than in other surface pelagic clusters, including sJB. Third, dinoflagellates were more relatively abundant in Hudson Bay than in James Bay, with the latter harbouring smaller taxa (Figure 6 and S2C). Dinoflagellates reached the highest relative abundance in oHB (Figure 5), which was mainly characterised by *Gyrodinium* and *Karlodinium*. *Biecheleria* characterised cHB, and *Gymnodinium* characterised James Bay. Summer surface waters transitioning from *Gyrodinium* in Hudson Bay to smaller *Gymnodinium* in James Bay aligns with Jacquemot et al. (2024), who noted that *Gymnodinium* prefers warmer and fresher waters. *Biecheleria* is likely common around the Belcher Islands, having been recovered there from sediment cores and traps in 2005-2006 (Heikkilä et al., 2014; 2016). This is the first observation of *Karlodinium* in the HBC. The *Karlodinium* taxon detected here could be the karlotoxin-producing *K. veneficum* (Liu et al., 2024), a harmful alga of concern, previously reported in the Canadian Arctic Ocean (Pućko et al., 2019; McKenzie et al., 2021) and recently observed to be relatively abundant in Disko Bay (Hoerstmann et al., 2025). *K. veneficum* is expected to form denser blooms in response to anticipated increases in temperature and light intensity (Coyne et al., 2021).

Cyanobacterial representation diverged between the surface pelagic waters of Hudson Bay and James Bay (Figure 6). James Bay was characterised by *Cyanobium* PCC-6307. Specifically, in southern James Bay, *Cyanobium* PCC-6307 was primarily represented by the estuarine-associated ASV_210, which occasionally exceeded 10% relative abundance (Figure S2D). *Cyanobium* is frequently reported to be relatively abundant in Hudson Bay rivers and their estuaries during summer, such as the Churchill, Nelson, and Great Whale Rivers (Blais et al., 2022; Morency et al., 2022). The strong presence of *Cyanobium* in James Bay reinforces that the bay, particularly its southern region, functions as an estuary. In contrast, Hudson Bay was characterised by *Synechococcus* CC9902. This study provides the first record of a relatively high abundance of *Synechococcus* within the HBC. Specifically, in 2023, the warm waters around the Belcher Islands exhibited a high relative abundance of *Synechococcus* CC9902, which was almost exclusively represented by the marine-associated ASV_17 and occasionally exceeded 10% prokaryotic relative abundance (Figure S2D). Previously, *Synechococcus* was detected in the HBC only within the Great Whale River, where it occurred at a low relative abundance of <0.5% (Blais et al., 2022). While Flombaum et al. (2013) projected a low presence of *Synechococcus* in Hudson Bay, those early models were understandably constrained by historically sparse regional sampling and a primary focus on temperature as an explanatory variable. However, Paulsen et al. (2016) and Doré et al. (2022) recently identified distinct high-latitude niches for *Synechococcus*, with Clade IV occupying the North Atlantic and Clade I favouring colder, saline Arctic waters, including the North Sea. While the peak relative abundance of ASV_17 occurred around a salinity of 27.5, suggesting affiliation with Clade I rather than Clade IV, Doré et al. (2022) noted that differences in salinity or temperature are insufficient to fully explain the distribution patterns of *Synechococcus*. Indeed, Celepli et al. (2017) documented that *Synechococcus* assemblages along the marine west coast of the Baltic Sea (salinity >20) were composed almost exclusively of marine strains most closely related to CC9311 (Clade I), CC9902 (Clade IV), and BL107 (Clade IV).

The transition from estuarine-associated *Cyanobium* PCC-6307 in James Bay to marine-associated *Synechococcus* CC9902 in Hudson Bay at a salinity of approximately 25 (Figure S2D) is very similar to the transition from *Cyanobium* PCC-7001 in the Baltic Sea to *Synechococcus* CC9311 in the North Sea at a salinity of approximately 15 (Celepli et al., 2017). Freshwater and estuarine *Cyanobium* strains (e.g., PCC7001 and PCC6307) harbour only phycocyanin (pigment type I), an adaptation to turbid, red-shifted conditions (Six et al., 2007; Larsson et al., 2014). In contrast, marine *Synechococcus* strains (e.g., CC9902 and CC9311) additionally possess phycoerythrin genes (pigment type III), which is an adaptation to clear, blue-shifted pelagic waters. Since the cyanobacterial pigment types have distinct spectral signatures, they can be used in remote sensing to trace changes in pelagic picocyanobacterial communities, serving as functional bioindicators of water-mass properties and ecosystem change (Mattei et al., 2026).

Measurements of fChl-a and beam attenuation (Figure 3B; Table S1), complemented by the preliminary findings for the R/V *William Kennedy* expeditions (Mundy et al., 2026), indicated that phytoplankton blooms occurred in the surface waters around the Belcher Islands in both 2022 and 2023 and near the Moose River in 2023. For Stn 8 sampled in 2022, Mundy et al. (2026) reported very high PP (3650 mg C m^-2^ d^-1^) and a surface pCO_2_ below atmospheric concentrations, while our results showed high relative abundance of *Chaetoceros* (Figure S2A). However, fChl-a at Stn 8 was below 1 mg m^-3^ at all depths (Figure 3B, BI22). It is likely that a strong, active *Chaetoceros* bloom occurred in the cold waters northwest of the Belcher Islands but was diluted by a relatively deep mixed layer created by strong tidal mixing (Mundy et al., 2026). In the warm waters around the Belcher Islands sampled in 2023, both high PP (Mundy et al., 2026) and elevated fChl-a (Figure 3B, >3.8 mg m^-3^ in cHB) were observed. Moreover, the elevated PP around the Belcher Islands coincided well with the elevated relative abundance of *Gyrodinium*, *Minidiscus*, and *Arcocellulus* in surface waters (Figures 5 and S2C). This observation suggests that in the warm waters around the Belcher Islands, there was a bloom of *Gyrodinium*, which, depending on the specific taxon, directly drove primary production or grazed on the small diatoms *Minidiscus* and *Arcocellulus*. *Gyrodinium* (ASV_635) also likely contributed to the vertical carbon export east of the islands (Stn 44), where it exceeded 50% relative abundance in subsurface waters (Figure 6). Finally, the waters near the Moose River sampled in 2023 exhibited the highest beam attenuation among all samples, which suggests a high concentration of suspended particulate matter in the water. This observation could reflect sediment resuspension caused by river discharge into the shallow coastal zone (bottom depth 6.4-7.6 m Stns 41-43) and/or an algal bloom. Indeed, fChl-a near the Moose River was also exceptionally high, reaching ∼7.5 mg m^-3^, far exceeding the median surface (1-6 m depth) value of ∼3.0 mg m^-3^ (n = 33) and paralleling the bloom peak recorded by a mooring in southern James Bay in 2022 (∼7.0 mg m^-3^; Mundy et al., 2026). Hence, a bloom of *Picochlorum* likely occurred near the Moose River in 2023 (Figure S2B). Indeed, the overwhelming presence of *Picochlorum* (ASV_680) in sJB likely resulted in the low eukaryotic diversity (Figure 5).

#### 4.1.2. Heterotrophic eukaryotes

In surface pelagic waters, differences in primary producers and water properties caused the divergence between heterotrophic eukaryotes of Hudson Bay and James Bay. Among grazers, the shift from high relative abundance of *Calanus* in Hudson Bay to that of *Pseudocalanus* in James Bay reflected an ecological transition from marine Arctic waters to river-influenced subarctic waters. The Arctic-adapted *Calanus glacialis* was strongly linked to colder water masses and elevated phytoplankton biomass in the HBC, where its presence in eastern Hudson Bay was driven by counterclockwise circulation advecting Arctic water and fauna from the Canadian Arctic Archipelago (Rochet and Grainger, 1988; Harvey et al., 2001; Schembri et al., 2023). It is likely that BI22 had an energy-efficient Arctic food web, where *Calanus* was grazing on the bloom of large *Chaetoceros* diatoms (Kohlbach et al., 2023). On the other hand, the river-influenced, subarctic waters of James Bay harboured the small copepod *Pseudocalanus*, which was determined by zooplankton sampling to dominate the copepod community in the bay (Mundy et al., 2026). *Pseudocalanus* was previously found to thrive in warm, fresh coastal zones, increasing in abundance from spring through its late summer reproductive period (Rochet and Grainger, 1988; Schembri et al., 2023). Among flagellates, Hudson Bay was characterised by *Cyrtostrombidium* in the cold water of 2022, and *Calliacantha* – in the warm waters of 2023, both of which were reported in Hudson Bay previously (Heikkilä et al., 2016; Jacquemot et al., 2022). On the other hand, James Bay was characterised *Ebria* and *Mesodinium. Ebria* is a coastal heterotrophic protist that feeds on diatoms and dinoflagellates (Hargraves, 2002) that was recently reported in the Great Whale River estuary in July (Jacquemot et al., 2021; Yurkowski et al., 2023). *Mesodinium* (ASV_668) characterizing James Bay may potentially be *M. rubrum*, a potentially harmful alga that retains plastids of consumed *Plagioselmis* (Stoecker and Lavrentyev, 2018). Recently, *M. rubrum* was reported as dominant in the Great Whale River estuary in July-August (Jacquemot et al., 2021; Blais et al., 2022). While it is possible that *Mesodinium* (ASV_668) could prey on *Plagioselmis* (ASV_725) – as both coexist in the warm waters around the Belcher Islands in 2023 and in James Bay – we did not find a statistically significant affiliation between the two. Finally, among Syndiniales, different DG-I-Clade-4 bioindicators characterised the surface waters of Hudson Bay and James Bay: ASV_954 in oHB and ASV_659 in sJB. Syndiniales are parasitic dinoflagellates that usually infect other dinoflagellates, ciliates, and crustaceans (Egge et al., 2025). Previously, DG-I, including Clade 4, was reported to be abundant offshore of the Great Whale River (Jacquemot et al., 2021) and in both surface and subsurface waters north of Svalbard in summer (Egge et al., 2025). Since Syndiniales succession follows that of their hosts, the bioindicator DG-I-Clade-4 likely infected *Gyrodinium* in oHB and *Gymnodinium* in sJB, respectively.

#### 4.1.3. Heterotrophic prokaryotes

In surface pelagic waters, differences in primary producers and water properties also caused a divergence between the heterotrophic prokaryotes of Hudson Bay and James Bay. Hudson Bay was mainly represented by members of Proteobacteria, Bacteroidota, and Verrucomicrobiota, which likely remineralized diatom-and dinoflagellate-derived organic matter in cold marine waters (Figures 4 and 5). RS62 marine group characterised the dinoflagellate-dominated cluster oHB, where it likely participated in the remineralization of *Gyrodinium*- and *Karlodinium*-derived organic matter, given its known positive response to phytoplankton-derived organic matter in the Baltic Sea (Elovaara et al., 2021). Interestingly, Verrucomicrobiota were also most abundant in oHB, where they were mainly represented by the particle-attached genus *Roseibacillus*, which is known to utilize complex organic substrates generated by phytoplankton blooms in the late Arctic summer (Delpech et al., 2021). *Sulfitobacter* and *Polaribacter* characterised the cold waters of BI22, where they likely utilized *Chaetoceros* exudates, as previously demonstrated by growth experiments with bacterial isolates from the Canadian Arctic Ocean (Tisserand et al., 2020). In contrast, the warm waters of cHB were characterised by *Formosa* and Pelagibacterales Clade Ia. *Formosa* likely utilized organic matter derived from the small polar diatoms *Minidiscus* and *Arcocellulus*, a link supported by an experiment involving a North Sea *Formosa* isolate (Unfried et al., 2018). *Formosa* also likely utilized organic matter derived from *Synechococcus* CC9902, given their co-occurance. Hence, while Flavobacteriaceae are well known to be abundant during phytoplankton blooms (Buchan et al., 2014), the Belcher Islands waters were occupied by *Polaribacter* in 2022, and *Formosa* in 2023, potentially reflecting substrate niche partitioning (Teeling et al., 2012) and/or temperature preference. Finally, Pelagibacterales Clade Ia is typically associated with stratified, post-bloom conditions, but certain cosmopolitan ecotypes can dominate less oligotrophic coastal environments (Delpech et al., 2021) and respond positively to tDOM (Lindh et al., 2015). Hence, Clade Ia could have benefited from exudates derived from the polar diatoms and dinoflagellates relatively abundant in cHB (Figure 6) and some of the tDOM transported by the southern Hudson Bay rivers and James Bay outflow towards cHB (median fCDOM 11.3 mg m^-3^; Figure 3B; Table S1).

James Bay was characterized by members of Actinomycetota and Planctomycetota, which were likely heterotrophic bacteria remineralizing green-algal, cyanobacterial, and terrestrial organic matter. Overall, sJB had a higher quantity of both autochthonous and allochthonous organic matter than nJB. As was mentioned above, there was likely a bloom of *Picochlorum* near the Moose River, and James Bay-specific *Cyanobium PCC-6307* was higher in sJB. Furthermore, while fCDOM was elevated near the Maquatua River (Stn 25) and near the Moose River (Stns 41-43), Stn 25 lay only 2 km offshore, but Stns 41-43 were 13-14 km from the coast. This suggests a much larger input of tDOM from the Moose River, as its signal persisted far from the river mouth. In support of this, southern James Bay was recently reported to contain ∼1.5 times more dissolved organic carbon than northern James Bay, which was attributed to the heavily regulated La Grande River delivering far less tDOM to northern James Bay than the unregulated Moose River to southern James Bay (Forget et al., 2025). Finally, James Bay was reported as net heterotrophic, with the highest air-sea CO_2_ flux occurring near the Moose River, suggesting active microbial degradation of organic matter (Mundy et al., 2026).

Several key taxa were identified as the most likely drivers of the net heterotrophy in James Bay. First, the NS3a marine group – a Flavobacteriaceae lineage distinct from *Polaribacter* and *Formosa* in Hudson Bay – characterised James Bay. Similar to other Flavobacteriaceae, NS3a marine group was previously observed to peak in response to phytoplankton blooms in the North and Baltic Seas (Teeling et al., 2016; Camarena-Gómez et al., 2018; von Scheibner et al., 2018), and its presence in James Bay was likely linked to utilization of *Picochlorum* and *Cyanobium* exudates. Second, both Sporichthyaceae and PeM15 belong to Actinomycetota, while *Fuerstia* belongs to Planctomycetota. Sporichthyaceae was previously reported in high relative abundance within the Great Whale River and its plume in summer, where it contributed substantially to Actinomycetota (Blais et al., 2022). Likewise, Planctomycetota was found to be associated with particles in the Great Whale River and its plume (Blais et al., 2022). *Fuerstia* in particular was previously identified in the Wadden Sea, an intertidal zone of the North Sea (Kohn et al., 2016). Given these riverine and coastal associations, it is likely that Sporichthyaceae and *Fuerstia* utilize tDOM in James Bay. Finally, *Pseudorhodobacter* was previously found in the Baltic Sea, where it was associated with macrophytes and suspected to degrade complex organic compounds such as lignin and polycyclic aromatic hydrocarbons (Ramasamy et al., 2025). Similarly, in eastern James Bay, the eelgrass collapse is driving the expansion of macroalgal mats and the release of sediment and organic matter, providing a substrate for *Pseudorhodobacter* (Kuzyk et al., 2026).

Different SAR86 bioindicators, ASV_18 and ASV_179, co-occurred with cyanobacteria in Hudson Bay and James Bay, respectively (Figure 6). Overall, members of the SAR86 clade are capable of utilizing cyanobacteria-derived organic matter, including sulfoquinovosyl diacylglycerol within their cell membranes (Dupont et al., 2012). Moreover, beyond this heterotrophic relationship, catalase-possessing SAR86 is also capable of protecting sensitive picocyanobacteria from oxidative stress (Braakman et al., 2017). The divergence of SAR86 clade taxa between Hudson Bay and James Bay could be driven by ASV_179 being specifically adapted to utilize the tDOM present in abundance in James Bay. Indeed, in incubation experiments performed using brackish waters from the Bothnian Sea and the Chukchi Sea, SAR86 clade was shown to increase and decrease, respectively, in response to higher tDOM concentrations (Lindh et al., 2015; Sipler et al., 2017).

### 4.2. River-estuary continuum

#### 4.2.1. Eukaryotic plankton

Eukaryotic communities of the Maquatua River (MR) flowing into James Bay and the Churchill and Winisk River estuaries (EST) located in Hudson Bay behaved differently from each other. Overall, the eukaryotic community of MR was very different from that in other clusters. MR had an order of magnitude lower relative abundance of dinoflagellates (3.5%) than other clusters and an exceptionally high relative abundance of fungi and telonemia (Figure 5). The bioindicator analysis of eukaryotes revealed that the MR taxa were restricted to the river, likely due to the near-zero salinity (Figure 6). Specifically, MR was represented by the photosynthetic algae *Choricystis* and *Aulacoseira*, the heterotrophic protists Telonemia-Group-2 and *Mesodinium*, and the fungus *Penicillium*, whose river-specific representatives were absent from or showed negligible presence in other clusters. *Choricystis* and *Penicillium* were previously reported in a permafrost thaw lake located near Kuujjuarapik in August (Przytulska et al., 2016). *Choricystis* also inhabits high-latitude freshwater and river-influenced marine environments, such as Lake Baikal and the Kara Sea (Kulakova et al., 2019; Belevich et al., 2021). Based on a recent mesocosm experiment using Gennarbyviken Reservoir freshwater, a high relative abundance of telonemia in the Maquatua River could signal enhanced nutrient cycling and primary productivity in response to warming, which ultimately increases food availability within the food web (Hall et al., 2025). The river-specific *Mesodinium* taxon is unlikely to be the potentially harmful *M. rubrum* suspected to occupy James Bay because its prey, *Plagioselmis* (exclusively ASV_725), was absent in MR (Figure 6).

In contrast, EST acted as a transitional environment between Hudson Bay and James Bay. EST clustered together with cHB and nJB/sJB along the primary axis of the dbRDA (Figure 4). The wide salinity range of EST samples (15.3-27.5) also suggested some tolerance to salinity changes (Table S1). Compared to other clusters, EST had an exceptionally high relative abundance of non-diatom Ochrophyta and Cryptophyta, with green algae and non-parasitic dinoflagellates also being relatively abundant (Figure 5). Yet, all EST bioindicators consistently appeared in other clusters, mainly cHB and nJB (Figure 6). Specifically, EST was characterised by *Aureococcus*, which was also relatively abundant in cHB, as well as *Plagioselmis*, *Micromonas*, and *Heterocapsa* (likely the mixotrophic dinoflagellate *H. rotundata*), which maintained elevated relative abundances in both cHB and nJB. Interestingly, despite the highest relative abundance of photosynthetic algae occurring in the Winisk River estuary, the Churchill River estuary (Stn 30) exhibited a distinct algal bloom signal (fChl-a 8.6 mg m^-3^). Echoing our findings, *Plagioselmis* and *Micromonas* occupied the Great Whale River plume (Blais et al., 2022) in August, and *H. rotundata* was relatively abundant in river-influenced eastern Hudson Bay in June-July (Amiraux et al., 2022; Jacquemot et al., 2024). To our knowledge, this was the first time *Aureococcus* was reported in Hudson Bay or James Bay. *Aureococcus* consists of the single species *A. anophagefferens*, which is a potentially harmful alga reported in the Canadian Arctic Ocean (Schiffrine et al., 2024). *A. anophagefferens* can create coastal brown tides that harm local flora and fauna (Gernez et al., 2023), and it was recently documented to be increasing in relative abundance in Disko Bay between 2018 and 2023 (Hoerstmann et al., 2025).

#### 4.2.2. Prokaryotic plankton

MR and EST harboured similar prokaryotic communities. Hierarchical clustering grouped MR and EST together (Figure 2), and several relatively abundant prokaryotic bioindicators were shared between the clusters. The similarity in riverine and estuarine prokaryotic communities is consistent with the mass effects paradigm (Leibold et al., 2004). This framework posits that sustained, high dispersal from a source habitat can maintain populations in a sink habitat, even where local conditions, such as a drastic salinity gradient, are unfavorable. Stadler and del Giorgio (2022) demonstrated these mass effects along the terrestrial-aquatic continuum of the La Romaine River in northeastern Canada. They showed that the downstream dispersal of terrestrial and riverine bacteria leaves a historical imprint in the estuary, where these transported organisms provide a major contribution to the estuarine bacterial community despite the vast majority being unreactive (detected through DNA, but not RNA). Parallel dynamics have been documented across other northern systems, including the Churchill River estuary, where river-derived, potentially halotolerant taxa constituted the majority of the bacterioplankton (Morency et al., 2022), and the Great Whale River plume, where a core freshwater microbiome persisted into the estuarine mixing zone (Blais et al., 2022). Complementing these past findings, the results in this study suggest that this fluvial carry-over is a widespread characteristic of river-estuary continuums across the broader Canadian Shield, where riverine taxa are rapidly transported to coastal waters across flat land underlain by impermeable bedrock. In this study, unidentified Sporichthyaceae as well as the *γ*-proteobacteria *Polynucleobacter* and *Candidatus* Methylopumilus characterised the Maquatua River but also persisted in the estuaries. These taxa and their distribution in river-estuary continuums match that for the Churchill, Nelson, Hayes, and Great Whale Rivers in summer (Blais et al., 2022; Morency et al., 2022). Both Sporichthyaceae and *Polynucleobacter* were also recently reported to be common in Svalbard lakes, accounting for up to 55% and 18% in relative abundance, respectively (Rappazzo et al., 2026). *Polynucleobacter* thrives on humic matter and phenol, core components of terrestrial dissolved organic matter (Hutalle-Schmelzer et al., 2010), while *Candidatus* Methylopumilus utilizes single-carbon substrates of cyanobacterial origin (Salcher et al., 2015). Finally, mass effects likely drove the elevated prokaryotic diversity in the river-influenced clusters (MR, EST, sJB, and dJB), with diversity declining northward and seaward (Figure 5).

Beyond the mass effects explanation, the similarity in prokaryotic communities between the riverine and estuarine environments of SHJB may also be explained by specific euryhaline prokaryotes capable of thriving in both environments. Despite drastically different salinities, both the Maquatua River and the Churchill and Winisk River estuaries were turbid, as evidenced by elevated beam attenuation. Furthermore, both the Maquatua River and the Churchill River estuary yielded the highest fCDOM values, pointing to high concentrations of terrestrial dissolved organic matter in both environments. Finally, dissolved lignin-phenol compositions were recently reported to be uniform among rivers flowing into both southern Hudson Bay and James Bay, suggesting similar terrestrial dissolved organic matter properties across the SHJB watershed (Forget et al., 2025). For example, Pelagibacterales Clade III, previously reported in high relative abundance in the Churchill, Nelson, and Hayes Rivers and their estuaries (Morency et al., 2022), was also prevalent in both the Maquatua River and the Churchill and Winisk River estuaries in this study. Pelagibacterales Clade III includes subclade IIIa, which is euryhaline and capable of growing across a wide range of temperatures (e.g., 12-35 °C for Arctic isolate IMCC9063; Oh et al., 2011; Lanclos et al., 2023). Some taxa of Pelagibacterales are also capable of utilizing both algal-derived and terrestrial dissolved organic matter (Lindh et al., 2015). *Cyanobium* PCC-6307 (ASV_221) appeared in MR but had a higher relative abundance in EST and even occurred in some offshore waters at a salinity of 29.2 (Figure S2D), which also suggests a high tolerance of this taxon to a wide salinity range spanning from fresh to marine waters.

### 4.3. Subsurface waters

The plankton communities of subsurface waters around the Belcher Islands (dBI) and in James Bay (dJB) were more similar to each other than to those in corresponding surface waters (Figure 2). dBI and dJB did not differ significantly in their physical parameters and were both deeper and colder than surface waters (Table S1), which according to dbRDA drove their planktonic communities to be distinct from those at the surface (Figure 4). Indeed, compared to surface waters, both dBI and dJB had an exceptionally high relative abundance of non-parasitic dinoflagellates and archaea. Nevertheless, despite surface and subsurface plankton communities being distinct from each other, the bioindicator analysis suggested a direct link between them, implying that subsurface communities were a combination of surface-derived and resident subsurface microorganisms, strongly influenced by the nature of sinking organic matter (Figure 6). Around the Belcher Islands, the bloom of *Gyrodinium* (ASV_635) in cHB was likely vertically transported to dBI. Interestingly, ASV_635 was also present in high relative abundance throughout the water column east of the Belcher Islands in 2022 (Stn 9), where there was deep mixed layer. *Gyrodinium* is a large heterotrophic dinoflagellate that can increase in size and become “fecal pellet-like” as it grazes on bloom-forming phytoplankton, causing it to readily sink and contribute to carbon export before being consumed by grazers (Matsuda et al., 2025). Nitrincolaceae co-occured with *Gyrodinium* in dBI. Given its strong presence in cHB and reports of its high relative abundance in surface waters of the summer Barents Sea during phytoplankton blooms (Thiele et al., 2023), it was likely associated with the phytoplankton growing in the surface waters around the Belcher Islands in 2023 (e.g., *Minidiscus*, *Arcocellulus*, *Micromonas*, and *Aureococcus*) and consumed by *Gyrodinium* as the dinoflagellates sank to depth. Given their high relative abundances in cHB that increased in dBI, *Pseudohongiella* and the SUP05 cluster were also likely associated with the sinking organic matter. *Pseudohongiella* was previously found to be associated with phytoplankton blooms in central Hudson Bay (Jacquemot et al., 2022), and the SUP05 cluster was relatively abundant in the deep northern Barents Sea (Thiele et al., 2023). In contrast, dJB was characterized by high relative abundances of not only non-parasitic dinoflagellates and archaea, but also Syndiniales, fungi, and Actinomycetota. Syndiniales from DG-I and DG-II were relatively abundant in dJB, likely infecting dinoflagellates (e.g., *Prorocentrum*), a pattern also observed in deep central Hudson Bay (Jacquemot et al., 2022). Given the relative abundance distribution of the Actinomycetota Sva0996 marine group, it was likely associated with the *Picochlorum* bloom in nJB/sJB and vertically transported to dJB with sinking aggregates of the green alga. Conversely, *Pseudomonas* and the archaeon *Candidatus* Nitrosopumilus appeared to be resident subsurface heterotrophs and chemotrophs, as they showed a weak presence in surface James Bay waters. *Sva0996* and the potentially ammonia-oxidizing archaeon *Candidatus* Nitrosopumilus were previously reported to be associated with sinking particles in deep central Hudson Bay (Jacquemot et al., 2022), while *Pseudomonas* was found to be associated with sinking particles in Baffin Bay (Amiraux et al., 2021).

## 5. Conclusion and future studies

This study offers new insights into the plankton composition of the Hudson Bay Complex (HBC) by presenting the first comprehensive environmental DNA survey of microorganisms in southern Hudson Bay and James Bay (SHJB). Surface pelagic communities shifted abruptly between marine-like Hudson Bay and river-influenced James Bay at a salinity of ∼24.5. Primary production shifted from large *Chaetoceros* diatoms in the cold (∼4 °C), well-mixed waters around the Belcher Islands in 2022 to smaller diatoms in the warmer (∼8 °C) coastal and Belcher Islands waters in 2023. James Bay was dominated by even smaller phototrophs – green algae (*Picochlorum*) and cyanobacteria (*Cyanobium*) – which had a stronger presence near the Moose River. Likewise, dinoflagellates and grazers also shifted in composition, with smaller taxa populating James Bay: Hudson Bay featured large *Gyrodinium* dinoflagellates and *Calanus* copepods, while James Bay favored *Gymnodinium* dinoflagellates and *Pseudocalanus* copepods, as well as small Cercozoa and Ciliophora protists. Finally, water properties and the nature and amount of organic matter shaped the heterotrophic bacteria. Based on the optical properties, fresher and warmer surface waters harboured more organic matter, derived from both primary production and rivers. Hudson Bay was mainly represented by Proteobacteria (RS62 marine group, *Sulfitobacter* and Pelagibacterales Clade Ia), Bacteroidota (*Polaribacter* and *Formosa*), and Verrucomicrobiota (*Roseibacillus*), which likely utilized diatom- and dinoflagellate-derived organic matter. On the other hand, James Bay was mainly represented by Actinomycetota (Sporichthyaceae and PeM15) and Planctomycetota (*Fuerstia*), which likely utilized *Cyanobium*-derived organic matter and tDOM. Some heterotrophic bacteria were cosmopolitan (*Planktomarina*, *Kineobactrum*, and *Candidatus* Aquiluna) across all non-riverine clusters, representing a background microbiome against which future shifts in SHJB could be measured.

In the Churchill and Winisk estuaries, eukaryotic communities were transitional between the coastal waters of Hudson Bay and James Bay, while prokaryotic communities were highly diverse and similar to those in the Maquatua River – a pattern that could be explained by the mass effects of downstream dispersal, with some taxa being euryhaline. In the Maquatua River, eukaryotic plankton (*Choricystis* and *Penicillium*) were restricted by a near-zero salinity, while the majority of prokaryotic plankton (Sporichthyaceae and *Polynucleobacter*) were suspected to utilize the in-situ tDOM. The microbial communities in subsurface waters were shaped by a vertical carbon flux of sinking dinoflagellates and algal aggregates, mainly shifting from *Gyrodinium*-associated marine bacteria (Nitrincolaceae) around the Belcher Islands to *Picochlorum*-associated aggregates (Sva0996 marine group) and Syndiniales in James Bay.

Based on the findings in this study, as the surface waters of the HBC become increasingly river-influenced, primary productivity and prokaryotic diversity will likely increase. However, due to a concurrent reduction in organism size, vertical carbon flux and energy transfer to higher trophic levels will likely become less efficient, which, coupled with elevated tDOM inputs, will result in a more dominant microbial loop (Assmy et al., 2023; Kohlbach et al., 2023). Additionally, we likely detected some harmful algae, such as toxin-producing *Karlodinium veneficum* and the brown-tide-forming *Aureococcus anophagefferens*, which are expected to increase in abundance in warmer surface waters, threatening both ecosystem and human health (Coyne et al., 2021; Hoerstmann et al., 2025). Finally, given the sharp salinity threshold between the basins, some of the ecosystem changes described may happen abruptly upon the water properties reaching a certain tipping point (Fu et al., 2020).

The DNA-based taxonomic analysis described here can be paired with physical and biogeochemical measurements to trace and predict ecosystem changes in the HBC and the rest of the Arctic Ocean. In the HBC, temperature, salinity and depth act as significant predictors of plankton community variability (this study; Jacquemot et al., 2024). Remote sensing and autonomous platforms are currently used in the HBC to monitor biogeochemically relevant parameters, such as temperature, salinity, and CDOM (Basu et al., 2026; Dmitrenko et al., 2021). Developing biogeochemical models for Hudson Bay (Deschepper et al., 2023) and the rest of the Canadian Arctic Ocean (Benoît-Gagné et al., 2024) can be improved by coupling biogeochemically relevant parameters with DNA-based taxonomic surveys to enable taxonomically resolved predictions of productivity, carbon flux, and ecosystem health. Additionally, this study provides the first record of a strong presence of a marine *Synechococcus* CC9902 strain in Hudson Bay, which contrasts with *Cyanobium* PCC-6307 taxa dominating cyanobacterial community in the river-estuary continuum and James Bay. Since these two picocyanobacterial groups differ in their pigment signatures, they can be monitored via remote sensing to track changes in surface water properties of Hudson and James Bay (Mattei et al., 2026). Finally, 16S rRNA sequencing can be applied in biodegradation assays (e.g., Kazmiruk et al., 2021) to identify bacteria responsible for remineralizing the increasing input of permafrost-derived and terrestrial organic matter into Hudson and James Bay, which would improve carbon flux estimates and predictions across the land-ocean continuum (Capelle et al., 2020).

## Acknowledgements

We thank the captain and crew of the R/V *William Kennedy* and the Southern Hudson Bay and James Bay expedition team, led by C. J. Mundy, Z. Z. A. Kuzyk, and J. K. Ehn, for field support during the 2022 and 2023 expeditions (Mundy et al., 2026).

## Supplementary material

Figures S1–S2 and Tables S1–S4 are provided in the accompanying Supplementary Material file.

## References

AMAP. 2024. AMAP Arctic Climate Change Update 2024: Key Trends and Impacts. Tromsø, Norway: Arctic Monitoring and Assessment Programme (AMAP). DOI: 10.21352/fz60-s852.

Amaral-Zettler, LA, McCliment, EA, Ducklow, HW, Huse, SM. 2009. A method for studying protistan diversity using massively parallel sequencing of V9 hypervariable regions of small-subunit ribosomal RNA genes. PLoS ONE 4(7): e6372. DOI: 10.1371/journal.pone.0006372.

Amaral-Zettler, LA, Bauer, M, Berg-Lyons, D, Betley, J, Caporaso, JG, Ducklow, HW, Fierer, N, Fraser, L, Gilbert, JA, Gormley, N, Huntley, J, Huse, SM, Jansson, JK, Jarman, SN, Knight, R, Lauber, CL, McCliment, EA, Owens, SM, Smith, G, Thompson, L, Vestheim, H, Walters, WA. 2023. EMP 18S Illumina amplicon protocol v2. protocols.io. DOI: 10.17504/protocols.io.ewov1b6pgr24/v2.

Amiraux, R, Rontani, J-F, Armougom, F, Frouin, E, Babin, M, Artigue, L, Bonin, P. 2021. Bacterial diversity and lipid biomarkers in sea ice and sinking particulate organic material during the melt season in the Canadian Arctic. Elementa: Science of the Anthropocene 9(1): 040. DOI: 10.1525/elementa.2019.040.

Amiraux, R, Lavaud, J, Cameron-Bergeron, K, Matthes, LC, Peeken, I, Mundy, CJ, Babb, DG, Tremblay, J-E. 2022. Content in fatty acids and carotenoids in phytoplankton blooms during the seasonal sea ice retreat in Hudson Bay complex, Canada. Elementa: Science of the Anthropocene 10(1): 00106. DOI: 10.1525/elementa.2021.00106.

Anderson, DM, Fachon, E, Pickart, RS, Lin, P, Fischer, AD, Richlen, ML, Uva, V, Brosnahan, ML, McRaven, L, Bahr, F, Lefebvre, K, Grebmeier, JM, Danielson, SL, Lyu, Y, Fukai, Y. 2021. Evidence for massive and recurrent toxic blooms of *Alexandrium catenella* in the Alaskan Arctic. Proceedings of the National Academy of Sciences 118(41): e2107387118. DOI: 10.1073/pnas.2107387118.

Apprill, A, McNally, S, Parsons, R, Weber, L. 2015. Minor revision to V4 region SSU rRNA 806R gene primer greatly increases detection of SAR11 bacterioplankton. Aquatic Microbial Ecology 75(2): 129–137. DOI: 10.3354/ame01753.

Assmy, P, Kvernvik, AC, Hop, H, Hoppe, CJM, Chierici, M, David, DT, Duarte, P, Fransson, A, García, LM, Patuła, W, Kwasniewski, S, Maturilli, M, Pavlova, O, Tatarek, A, Wiktor, JM, Wold, A, Wolf, KKE, Bailey, A. 2023. Seasonal plankton dynamics in Kongsfjorden during two years of contrasting environmental conditions. Progress in Oceanography 213: 102996. DOI: 10.1016/j.pocean.2023.102996.

Baker, ME, King, RS. 2010. A new method for detecting and interpreting biodiversity and ecological community thresholds. Methods in Ecology and Evolution 1(1): 25–37. DOI: 10.1111/j.2041-210X.2009.00007.x.

Basu, A, Bélanger, S, Kuzyk, ZZA, Singh, RK, Mukherjee, S, Larouche, P, Guzzi, AC, Ausen, EL, McCullough, G, Ehn, JK. 2026. Satellite data trace seasonal freshwater dispersion in Hudson and James Bays. Geophysical Research Letters 53(2): e2025GL116442. DOI: 10.1029/2025GL116442.

Belevich, TA, Ilyash, LV, Milyutina, IA, Troitsky, AV, Sergeeva, VM. 2019. Picoplanktonic diatoms of Russian Arctic Seas revealed by metagenome analyze. Voprosy sovremennoi algologii 2(20): 105–110. DOI: 10.33624/2311-0147-2019-2(20)-105-110.

Belevich, TA, Milyutina, IA, Troitsky, AV. 2021. Seasonal variability of photosynthetic microbial eukaryotes (*<*3 µm) in the Kara Sea revealed by 18S rDNA metabarcoding of sediment trap fluxes. Plants 10(11): 2394. DOI: 10.3390/plants10112394.

Benoît-Gagné, M, Dutkiewicz, S, Deschepper, I, Dufresne, C, Dumont, D, Larouche, R, Mémery, L, Olivier, G, Maps, F. 2024. Exploring controls on the timing of the phytoplankton bloom in western Baffin Bay, Canadian Arctic. Elementa: Science of the Anthropocene 12(1): 00008. DOI: 10.1525/elementa.2024.00008.

Blais, MA, Matveev, A, Lovejoy, C, Vincent, WF. 2022. Size-fractionated microbiome structure in subarctic rivers and a coastal plume across DOC and salinity gradients. Frontiers in Microbiology 12: 760282. DOI: 10.3389/fmicb.2021.760282.

Braakman, R, Follows, MJ, Chisholm, SW. 2017. Metabolic evolution and the self-organization of ecosystems. Proceedings of the National Academy of Sciences 114(15): E3091–E3100. DOI: 10.1073/pnas.1619573114.

Buchan, A, LeCleir, GR, Gulvik, CA, González, JM. 2014. Master recyclers: Features and functions of bacteria associated with phytoplankton blooms. Nature Reviews Microbiology 12(10): 686–698. DOI: 10.1038/nrmicro3326.

Callahan, BJ, McMurdie, PJ, Rosen, MJ, Han, AW, Johnson, AJA, Holmes, SP. 2016. DADA2: High-resolution sample inference from Illumina amplicon data. Nature Methods 13(7): 581–583. DOI: 10.1038/nmeth.3869.

Camarena-Gómez, MT, Lipsewers, T, Piiparinen, J, Eronen-Rasimus, E, Perez-Quemalinos, D, Hoikkala, L, Sobrino, C, Spilling, K. 2018. Shifts in phytoplankton community structure modify bacterial production, abundance and community composition. Aquatic Microbial Ecology 81(2): 149–170. DOI: 10.3354/ame01868.

Capelle, DW, Kuzyk, ZZA, Papakyriakou, T, Guéguen, C, Miller, LA, Macdonald, RW. 2020. Effect of terrestrial organic matter on ocean acidification and CO_2_ flux in an Arctic shelf sea. Progress in Oceanography 185: 102319. DOI: 10.1016/j.pocean.2020.102319.

Caporaso, JG, Lauber, CL, Walters, WA, Berg-Lyons, D, Lozupone, CA, Turnbaugh, PJ, Fierer, N, Knight, R. 2011. Global patterns of 16S rRNA diversity at a depth of millions of sequences per sample. Proceedings of the National Academy of Sciences 108(Suppl. 1): 4516–4522. DOI: 10.1073/pnas.1000080107.

Caporaso, JG, Ackermann, G, Apprill, A, Bauer, M, Berg-Lyons, D, Betley, J, Fierer, N, Fraser, L, Fuhrman, JA, Gilbert, JA, Gormley, N, Humphrey, G, Huntley, J, Jansson, JK, Knight, R, Lauber, CL, Lozupone, CA, McNally, S, Needham, DM, Owens, SM, Parada, AE, Parsons, R, Smith, G, Thompson, LR, Thompson, L, Turnbaugh, PJ, Walters, WA, Weber, L. 2023. EMP 16S Illumina amplicon protocol v2. protocols.io. DOI: 10.17504/protocols.io.kqdg3dzzl25z/v2.

Celepli, N, Sundh, J, Ekman, M, Dupont, CL, Yooseph, S, Bergman, B, Ininbergs, K. 2017. Meta-omic analyses of Baltic Sea cyanobacteria: Diversity, community structure and salt acclimation. Environmental Microbiology 19(2): 673–686. DOI: 10.1111/1462-2920.13592.

Champagne, O, Arain, MA, Wang, S, Leduc, M, Russell, HAJ. 2021. Interdecadal variability of streamflow in the Hudson Bay Lowlands watersheds driven by atmospheric circulation. Journal of Hydrology: Regional Studies 36: 100868. DOI: 10.1016/j.ejrh.2021.100868.

Comeau, AM, Li, WKW, Tremblay, J-É, Carmack, EC, Lovejoy, C. 2011. Arctic Ocean microbial community structure before and after the 2007 record sea ice minimum. PLoS ONE 6(11): e27492. DOI: 10.1371/journal.pone.0027492.

Coyne, KJ, Salvitti, LR, Mangum, AM, Ozbay, G, Main, CR, Kouhanestani, ZM, Warner, ME. 2021. Interactive effects of light, CO_2_ and temperature on growth and resource partitioning by the mixotrophic dinoflagellate, *Karlodinium veneficum*. PLoS ONE 16(10): e0259161. DOI: 10.1371/journal.pone.0259161.

Cruaud, P, Vigneron, A, Fradette, MS, Charette, SJ, Rodriguez, MJ, Dorea, CC, Culley, AI. 2017. Open the Sterivex casing: An easy and effective way to improve DNA extraction yields. Limnology and Oceanography: Methods 15(12): 1015–1020. DOI: 10.1002/lom3.10221.

De Cáceres, M, Legendre, P. 2009. Associations between species and groups of sites: Indices and statistical inference. Ecology 90(12): 3566–3574. DOI: 10.1890/08-1823.1.

Delpech, L-M, Vonnahme, TR, McGovern, M, Gradinger, R, Præbel, K, Poste, AE. 2021. Terrestrial inputs shape coastal bacterial and archaeal communities in a high Arctic fjord (Isfjorden, Svalbard). Frontiers in Microbiology 12: 614634. DOI: 10.3389/fmicb.2021.614634.

Deschepper, I, Myers, PG, Lavoie, D, Papakyriakou, T, Maps, F. 2023. Understanding the physical forcings behind the biogeochemical productivity of the Hudson Bay Complex. Journal of Geophysical Research: Biogeosciences 128(6): e2022JG007294. DOI: 10.1029/2022JG007294.

Di Pane, J, Bourdaud, P, Horn, S, Duarte Moreno, H, Meunier, CL. 2024. Global change alters coastal plankton food webs by promoting the microbial loop: An inverse modelling and network analysis approach on a mesocosm experiment. Science of the Total Environment 921: 171272. DOI: 10.1016/j.scitotenv.2024.171272.

Dickey, MH. 2015. Status of eelgrass beds on the east coast of James Bay. Québec, QC: Environment Canada, Wildlife and Landscape Science Directorate. Available at https://publications.gc.ca/site/eng/9.898377/publication.html.

Dmitrenko, IA, Kirillov, SA, Babb, DG, Kuzyk, ZZA, Basu, A, Ehn, JK, Sydor, K, Barber, DG. 2021. Storm-driven hydrography of western Hudson Bay. Continental Shelf Research 227: 104525. DOI: 10.1016/j.csr.2021.104525.

Doré, H, Leconte, J, Guyet, U, Breton, S, Farrant, GK, Demory, D, Ratin, M, Hoebeke, M, Corre, E, Pitt, FD, Ostrowski, M, Scanlan, DJ, Partensky, F, Six, C, Garczarek, L. 2022. Global phylogeography of marine *Synechococcus* in coastal areas reveals strong community shifts. mSystems 7(6): e00656–22. DOI: 10.1128/msystems.00656-22.

Dufrêne, M, Legendre, P. 1997. Species assemblages and indicator species: The need for a flexible asymmetrical approach. Ecological Monographs 67(3): 345–366. DOI: 10.1890/0012-9615(1997)067[0345:SAAIST]2.0.CO;2.

Dupont, CL, Rusch, DB, Yooseph, S, Lombardo, MJ, Richter, RA, Valas, R, Novotny, M, Yee-Greenbaum, J, Selengut, JD, Haft, DH, Halpern, AL, Lasken, RS, Nealson, K, Friedman, R, Venter, JC. 2012. Genomic insights to SAR86, an abundant and uncultivated marine bacterial lineage. The ISME Journal 6(6): 1186– 1199. DOI: 10.1038/ismej.2011.189.

Eastwood, RA, Macdonald, RW, Ehn, JK, Heath, J, Arragutainaq, L, Myers, PG, Barber, DG, Kuzyk, ZA. 2020. Role of river runoff and sea ice brine rejection in controlling stratification throughout winter in southeast Hudson Bay. Estuaries and Coasts 43(4): 756–786. DOI: 10.1007/s12237-020-00698-0.

Egge, E, Vaulot, D, Larsen, A, Edvardsen, B. 2025. The community of marine alveolate parasites in the Atlantic inflow to the Arctic Ocean is structured by season, depth, and water mass. Arctic Science 11: 1–16. DOI: 10.1139/as-2024-0016.

Elovaara, S, Eronen-Rasimus, E, Asmala, E, Tamelander, T, Kaartokallio, H. 2021. Contrasting patterns of carbon cycling and dissolved organic matter processing in two phytoplankton–bacteria communities. Biogeosciences 18(24): 6589–6616. DOI: 10.5194/bg-18-6589-2021.

Flombaum, P, Gallegos, JL, Gordillo, RA, Rincón, J, Zabala, LL, Jiao, N, Karl, DM, Li, WKW, Lomas, MW, Veneziano, D, Vera, CS, Vrugt, JA, Martiny, AC. 2013. Present and future global distributions of the marine cyanobacteria *Prochlorococcus* and *Synechococcus*. Proceedings of the National Academy of Sciences 110(24): 9824–9829. DOI: 10.1073/pnas.1307701110.

Forget, JP, Kuzyk, ZZ, Mundy, CJ, Guéguen, C. 2025. Dissolved organic matter (DOM) and barium in James Bay: Distribution, sources, and climate change implications. Journal of Marine Systems 250: 104084. DOI: 10.1016/j.jmarsys.2025.104084.

Fu, W, Moore, JK, Primeau, FW, Lindsay, K, Randerson, JT. 2020. A growing freshwater lens in the Arctic Ocean with sustained climate warming disrupts marine ecosystem function. Journal of Geophysical Research: Biogeosciences 125(12): e2020JG005693. DOI: 10.1029/2020JG005693.

Gernez, P, Zoffoli, ML, Lacour, T, Hernández Farinas, T, Navarro, G, Caballero, I, Harmel, T. 2023. The many shades of red tides: Sentinel-2 optical types of highly-concentrated harmful algal blooms. Remote Sensing of Environment 287: 113486. DOI: 10.1016/j.rse.2023.113486.

Gough, WA, Leung, A. 2002. Nature and fate of Hudson Bay permafrost. Regional Environmental Change 2(4): 177–184. DOI: 10.1007/s10113-002-0048-x.

Granskog, MA, Kuzyk, ZZA, Azetsu-Scott, K, Macdonald, RW. 2011. Distributions of runoff, sea-ice melt and brine using *δ*^18^O and salinity data – A new view on freshwater cycling in Hudson Bay. Journal of Marine Systems 88(3): 362–374. DOI: 10.1016/j.jmarsys.2011.03.011.

Guillou, L, Bachar, D, Audic, S, Bass, D, Berney, C, Bittner, L, Boutte, C, Burgaud, G, De Vargas, C, Decelle, J, Del Campo, J, Dolan, JR, Dunthorn, M, Edvardsen, B, Holzmann, M, Kooistra, WHCF, Lara, E, Le Bescot, N, Logares, R, Mahé, F, Massana, R, Montresor, M, Morard, R, Not, F, Pawlowski, J, Probert, I, Sauvadet, AL, Siano, R, Stoeck, T, Vaulot, D, Zimmermann, P, Christen, R. 2013. The Protist Ribosomal Reference database (PR2): A catalog of unicellular eukaryote Small Sub-Unit rRNA sequences with curated taxonomy. Nucleic Acids Research 41(D1): D597–D604. DOI: 10.1093/nar/gks1160.

Hahsler, M, Nagar, A. 2024. rBLAST: R interface for the Basic Local Alignment Search Tool. Bioconductor. DOI: 10.18129/B9.bioc.rBLAST.

Hall, CAM, Henry, N, Canals, O, Consing, G, Rodríguez-Ezpeleta, N, Lewandowska, AM. 2025. DNA metabarcoding as a tool to study plankton responses to warming and salinity change in mesocosms. Ecology and Evolution 15(9): e72125. DOI: 10.1002/ece3.72125.

Hargraves, PE. 2002. The ebridian flagellates *Ebria* and *Hermesinum*. Plankton Biology and Ecology 49(1): 9–16.

Harris, LI, Richardson, K, Bona, KA, Davidson, SJ, Finkelstein, SA, Garneau, M, McLaughlin, J, Nwaishi, F, Olefeldt, D, Packalen, M, Roulet, NT, Southee, FM, Strack, M, Webster, KL, Wilkinson, SL, Ray, JC. 2022. The essential carbon service provided by northern peatlands. Frontiers in Ecology and the Environment 20(4): 222–230. DOI: 10.1002/fee.2437.

Harvey, M, Therriault, J-C, Simard, N. 2001. Hydrodynamic control of late summer species composition and abundance of zooplankton in Hudson Bay and Hudson Strait (Canada). Journal of Plankton Research 23(5): 481–499. DOI: 10.1093/plankt/23.5.481.

Heikkilä, M, Pospelova, V, Hochheim, KP, Kuzyk, ZZA, Stern, GA, Barber, DG, Macdonald, RW. 2014. Surface sediment dinoflagellate cysts from the Hudson Bay system and their relation to freshwater and nutrient cycling. Marine Micropaleontology 106: 79–109. DOI: 10.1016/j.marmicro.2013.12.002.

Heikkilä, M, Pospelova, V, Forest, A, Stern, GA, Fortier, L, Macdonald, RW. 2016. Dinoflagellate cyst production over an annual cycle in seasonally ice-covered Hudson Bay. Marine Micropaleontology 125: 1–24. DOI: 10.1016/j.marmicro.2016.02.005.

Hoerstmann, C, Hansen, PJ, Lundholm, N, Krock, B, John, U. 2025. Multi-year dynamics of harmful algae in Disko Bay, West Greenland. Harmful Algae 145: 102857. DOI: 10.1016/j.hal.2025.102857.

Husson, B, Bluhm, BA, Cyr, F, Danielson, SL, Eriksen, E, Fossheim, M, Geoffroy, M, Hopcroft, RR, Ingvaldsen, RB, Jørgensen, LL, Lovejoy, C, Meire, L, Mueter, F, Primicerio, R, Winding, M. 2024. Borealization impacts shelf ecosystems across the Arctic. Frontiers in Environmental Science 12: 1481420. DOI: 10.3389/fenvs.2024.1481420.

Hutalle-Schmelzer, KML, Zwirnmann, E, Krüger, A, Grossart, HP. 2010. Enrichment and cultivation of pelagic bacteria from a humic lake using phenol and humic matter additions. FEMS Microbiology Ecology 72(1): 58–73. DOI: 10.1111/j.1574-6941.2009.00831.x.

Jackson, VLN, Grevesse, T, Kilias, ES, Onda, DFL, Young, KF, Allen, MJ, Walsh, DA, Lovejoy, C, Monier, A. 2024. Vulnerability of Arctic Ocean microbial eukaryotes to sea ice loss. Scientific Reports 14(1): 28896. DOI: 10.1038/s41598-024-77821-9.

Jacquemot, L, Kalenitchenko, D, Matthes, LC, Vigneron, A, Mundy, CJ, Tremblay, J-É, Lovejoy, C. 2021. Protist communities along freshwater–marine transition zones in Hudson Bay (Canada). Elementa: Science of the Anthropocene 9(1): 00111. DOI: 10.1525/elementa.2021.00111.

Jacquemot, L, Vigneron, A, Tremblay, J-É, Lovejoy, C. 2022. Contrasting sea ice conditions shape microbial food webs in Hudson Bay (Canadian Arctic). ISME Communications 2(1): 104. DOI: 10.1038/s43705-022-00192-7.

Jacquemot, L, Tremblay, J-É, Morency, C, Lovejoy, C. 2024. Stratification and summer protist communities in the Arctic influenced coastal systems of Nunavik (Québec, Canada). Frontiers in Marine Science 11: 1321604. DOI: 10.3389/fmars.2024.1321604.

Kaczmarska, I, Lovejoy, C, Potvin, M, MacGillivary, M. 2009. Morphological and molecular characteristics of selected species of *Minidiscus* (Bacillariophyta, Thalassiosiraceae). European Journal of Phycology 44(4): 461–475. DOI: 10.1080/09670260902855873.

Kazmiruk, ZV, Capelle, DW, Kamula, CM, Rysgaard, S, Papakyriakou, T, Kuzyk, ZA. 2021. High biodegradability of riverine dissolved organic carbon in late winter in Hudson Bay, Canada. Elementa: Science of the Anthropocene 9(1): 00123. DOI: 10.1525/elementa.2020.00123.

Kellogg, CTE, McClelland, JW, Dunton, KH, Crump, BC. 2019. Strong seasonality in Arctic estuarine microbial food webs. Frontiers in Microbiology 10: 2628. DOI: 10.3389/fmicb.2019.02628.

Kohlbach, D, Goraguer, L, Bodur, YV, Müller, O, Amargant-Arumí, M, Blix, K, Bratbak, G, Chierici, M, Dąbrowska, AM, Dietrich, U, Edvardsen, B, García, LM, Gradinger, R, Hop, H, Jones, E, Lundesgaard, Ø, Olsen, LM, Reigstad, M, Saubrekka, K, Tatarek, A, Wiktor, JM, Wold, A, Assmy, P. 2023. Earlier sea-ice melt extends the oligotrophic summer period in the Barents Sea with low algal biomass and associated low vertical flux. Progress in Oceanography 213: 103018. DOI: 10.1016/j.pocean.2023.103018.

Kohn, T, Heuer, A, Jogler, M, Vollmers, J, Boedeker, C, Bunk, B, Rast, P, Borchert, D, Glöckner, I, Freese, HM, Klenk, HP, Overmann, J, Kaster, AK, Rohde, M, Wiegand, S, Jogler, C. 2016. *Fuerstia marisgermanicae* gen. nov., sp. nov., an unusual member of the phylum Planctomycetes from the German Wadden Sea. Frontiers in Microbiology 7: 2079. DOI: 10.3389/fmicb.2016.02079.

Kulakova, NV, Kashin, SA, Bukin, YS. 2019. The genetic diversity and phylogeny of green microalgae in the genus *Choricystis* (Trebouxiophyceae, Chlorophyta) in Lake Baikal. Limnology 21(1): 15–24. DOI: 10.1007/s10201-019-00587-x.

Kuzyk, ZZA, Leblanc, M, Ehn, J, Crawford, A, Peck, C, Guzzi, A, Stocking, M, Idrobo, CJ, Knight, NS, Davis, K, Dunn, M, Rabbitskin, E, Bélanger, S, Noisette, F, Gosselin, M, Fink-Mercier, C, Neumeier, U, Walch, D, De Melo, M, Del Giorgio, PA, Humphries, M, O’Connor, MI. 2026. Eelgrass ecosystem collapse and social-ecological regime shift driven by hydropower development and climate change. Nature Communications 17(1): 2917. DOI: 10.1038/s41467-026-69553-3.

Lanclos, VC, Rasmussen, AN, Kojima, CY, Cheng, C, Henson, MW, Faircloth, BC, Francis, CA, Thrash, JC. 2023. Ecophysiology and genomics of the brackish water adapted SAR11 subclade IIIa. The ISME Journal 17(4): 620–629. DOI: 10.1038/s41396-023-01376-2.

Larsson, J, Celepli, N, Ininbergs, K, Dupont, CL, Yooseph, S, Bergman, B, Ekman, M. 2014. Picocyanobacteria containing a novel pigment gene cluster dominate the brackish water Baltic Sea. The ISME Journal 8(9): 1892–1903. DOI: 10.1038/ismej.2014.35.

Leibold, MA, Holyoak, M, Mouquet, N, Amarasekare, P, Chase, JM, Hoopes, MF, Holt, RD, Shurin, JB, Law, R, Tilman, D, Loreau, M, Gonzalez, A. 2004. The metacommunity concept: A framework for multi-scale community ecology. Ecology Letters 7(7): 601–613. DOI: 10.1111/j.1461-0248.2004.00608.x.

Li, Y, Han, D, Rogers, CA, Finkelstein, SA, Hararuk, O, Waddington, JM, Barreto, C, McLaughlin, JW, Snider, J, Gonsamo, A. 2025. Peat depth and carbon storage of the Hudson Bay Lowlands, Canada. Geophysical Research Letters 52(2): e2024GL110679. DOI: 10.1029/2024GL110679.

Lindh, MV, Lefébure, R, Degerman, R, Lundin, D, Andersson, A, Pinhassi, J. 2015. Consequences of increased terrestrial dissolved organic matter and temperature on bacterioplankton community composition during a Baltic Sea mesocosm experiment. Ambio 44(Suppl. 3): 402–412. DOI: 10.1007/s13280-015-0659-3.

Liu, X, Bian, Z, Hu, S, Dickinson, CF, Benjamin, MM, Jia, J, Tian, Y, Place, A, Hanna, GS, Luesch, H, Croot, P, Reddy, MM, Thomas, OP, Hardiman, G, Puglisi, MP, Yang, M, Zhong, Z, Lemasters, JJ, Korte, JE, Waters, AL, Heltzel, CE, Williamson, RT, Strangman, WK, Valeriote, F, Tius, MA, DiTullio, GR, Ferreira, D, Alekseyenko, A, Wang, S, Hamann, MT, Wang, X. 2024. The chemistry of phytoplankton. Chemical Reviews 124(23): 13099–13177. DOI: 10.1021/acs.chemrev.4c00177.

Loutet, SJT, Sanger, A, Strong, K, Collins, RE, Mahmoudi, N. 2025. Microbial communities in the Hudson Strait amidst rapid environmental change. Canadian Journal of Microbiology 71: 1–9. DOI: 10.1139/cjm-2024-0154.

MacDonald, MK, Stadnyk, TA, Déry, SJ, Braun, M, Gustafsson, D, Isberg, K, Arheimer, B. 2018. Impacts of 1.5 and 2.0 °C warming on pan-Arctic river discharge into the Hudson Bay Complex through 2070. Geophysical Research Letters 45(15): 7561–7570. DOI: 10.1029/2018GL079147.

Matsuda, R, Sano, M, Tsuchiya, C, Takahashi, KD, Takao, S, Mizobata, K, Moteki, M, Makabe, R, Kurosawa, N. 2025. Spatial and temporal occurrence of two *Gyrodinium* species in the seasonal ice zone off Wilkes Land, Southern Ocean. Polar Science 43: 101169. DOI: 10.1016/j.polar.2025.101169.

Mattei, F, Hickman, AE, Uitz, J, Vellucci, V, Garczarek, L, Partensky, F, Dutkiewicz, S. 2026. Phytoplankton with flexible pigment content disadvantaged by projected future decrease in variability of the ocean light spectrum. Global Change Biology 32(1): e70671. DOI: 10.1111/gcb.70671.

McCullough, G, Kuzyk, ZZA, Ehn, J, Babb, DG, Ridenour, N, Myers, PG, Wong, K, Koenig, K, Sydor, K, Barber, D. 2019. Freshwater–marine interactions in the Greater Hudson Bay Marine Region, in Kuzyk, ZA, Candlish, LM, editors. From science to policy in the Greater Hudson Bay Marine Region: An Integrated Regional Impact Study (IRIS) of climate change and modernization. Québec City, Québec, Canada: ArcticNet. p. 113–133.

McKenzie, CH, Bates, SS, Martin, JL, Haigh, N, Howland, KL, Lewis, NI, Locke, A, Peña, A, Poulin, M, Rochon, A, Rourke, WA, Scarratt, MG, Starr, M, Wells, T. 2021. Three decades of Canadian marine harmful algal events: Phytoplankton and phycotoxins of concern to human and ecosystem health. Harmful Algae 102: 101852. DOI: 10.1016/j.hal.2020.101852.

Mordret, S, MacKinnon, J, Behnke, J, O’Leary, SJB, Chénard, C. 2024. Identification of phytoplankton isolates from the eastern Canadian waters using long-read sequencing. Journal of Plankton Research 46(6): 527–541. DOI: 10.1093/plankt/fbae043.

Morency, C, Jacquemot, L, Potvin, M, Lovejoy, C. 2022. A microbial perspective on the local influence of Arctic rivers and estuaries on Hudson Bay (Canada). Elementa: Science of the Anthropocene 10(1): 00009. DOI: 10.1525/elementa.2021.00009.

Mundy, CJ, Ausen, E, Ahrenholtz, H, Yezhova, K, Kirillov, S, Kuzyk, Z, Decker, N, Kazmiruk, Z, Collins, E, Vachon, N, Niemi, A, Long, A, Bedington, M, Jones, E, Ehn, J, Guzzi, A, Guillaume, M, Bouchard, P, Capelle, D, Papakyriakou, T, Plouffe, J, Yurkowski, D, Woodard, K, Marcoux, M. 2026. Southern Hudson Bay – James Bay Knowledge Synthesis: Summary of Recent Oceanographic Research in James Bay and Belcher Islands. Centre for Earth Observation Science (CEOS).

Oh, HM, Kang, I, Lee, K, Jang, Y, Lim, SI, Cho, JC. 2011. Complete genome sequence of strain IMCC9063, belonging to SAR11 subgroup 3, isolated from the Arctic Ocean. Journal of Bacteriology 193(13): 3379– 3380. DOI: 10.1128/JB.05033-11.

Oksanen, J, Simpson, GL, Blanchet, FG, Kindt, R, Legendre, P, Minchin, PR, O’Hara, RB, Solymos, P, Stevens, MHH, Szoecs, E, Wagner, H, Bedward, M, Bolker, B, Borcard, D, Carvalho, G, De Caceres, M, Durand, S, Evangelista, HBA, Hannigan, G, Hill, MO, Lahti, L, Martino, C, Ouellette, MH, Ribeiro Cunha, E, Smith, T, Stier, A, Ter Braak, CJF, Weedon, J. 2026. vegan: Community Ecology Package. Version 2.7.5. DOI: 10.32614/CRAN.package.vegan.

Paulsen, ML, Doré, H, Garczarek, L, Seuthe, L, Müller, O, Sandaa, RA, Bratbak, G, Larsen, A. 2016. *Synechococcus* in the Atlantic Gateway to the Arctic Ocean. Frontiers in Marine Science 3: 191. DOI: 10.3389/fmars.2016.00191.

Peck, GS. 1976. James Bay Oceanographic Data Report – Volume 1: Winter 1975–1976.

Peralta-Ferriz, C, Woodgate, RA. 2015. Seasonal and interannual variability of pan-Arctic surface mixed layer properties from 1979 to 2012 from hydrographic data, and the dominance of stratification for multiyear mixed layer depth shoaling. Progress in Oceanography 134: 19–53. DOI: 10.1016/j.pocean.2014.12.005.

Przytulska, A, Comte, J, Crevecoeur, S, Lovejoy, C, Laurion, I, Vincent, WF. 2016. Phototrophic pigment diversity and picophytoplankton in permafrost thaw lakes. Biogeosciences 13(1): 13–26. DOI: 10.5194/bg-13-13-2016.

Pućko, M, Dionne, K, Michel, C. 2019. Occurrence of toxin-producing marine algae in the Canadian Arctic and adjacent waters. Ottawa: Fisheries and Oceans Canada. Canadian Manuscript Report of Fisheries and Aquatic Sciences 3180. Available at https://publications.gc.ca/site/eng/9.874970/publication.html.

Quast, C, Pruesse, E, Yilmaz, P, Gerken, J, Schweer, T, Yarza, P, Peplies, J, Glöckner, FO. 2013. The SILVA ribosomal RNA gene database project: Improved data processing and web-based tools. Nucleic Acids Research 41(D1): D590–D596. DOI: 10.1093/nar/gks1219.

Ramasamy, KP, Vass, M, Berglund, J, Saarinen, A, Andersson, A. 2025. Perfoliate pondweed meadows in northern coastal areas: Reservoirs of diverse bacteria with pathogenic traits and complex carbon metabolism. Environmental Microbiology Reports 17(5): e70180. DOI: 10.1111/1758-2229.70180.

Rantanen, M, Karpechko, AY, Lipponen, A, Nordling, K, Hyvärinen, O, Ruosteenoja, K, Vihma, T, Laaksonen, A. 2022. The Arctic has warmed nearly four times faster than the globe since 1979. Communications Earth & Environment 3(1): 168. DOI: 10.1038/s43247-022-00498-3.

Rappazzo, AC, Lo Giudice, A, Rizzo, C, Giannarelli, S, Ghezzi, L, Azzaro, M, Cairns, WRL, Papale, M. 2026. Bacterial community and chemical contamination in high-latitude lake waters: A comparative study from the Svalbard and Antarctic Peninsula regions. Environmental Research 295: 123915. DOI: 10.1016/j.envres.2026.123915.

Ribeiro, CG, dos Santos, AL, Gourvil, P, Le Gall, F, Marie, D, Tragin, M, Probert, I, Vaulot, D. 2020. Culturable diversity of Arctic phytoplankton during pack ice melting. Elementa: Science of the Anthropocene 8: 6. DOI: 10.1525/elementa.401.

Robicheau, BM, Tolman, J, Bertrand, EM, LaRoche, J. 2022. Highly-resolved interannual phytoplankton community dynamics of the coastal Northwest Atlantic. ISME Communications 2(1): 38. DOI: 10.1038/s43705-022-00119-2.

Rochet, M, Grainger, EH. 1988. Community structure of zooplankton in eastern Hudson Bay. Canadian Journal of Zoology 66(7): 1626–1630. DOI: 10.1139/z88-237.

Royo-Llonch, M, Sánchez, P, Ruiz-González, C, Salazar, G, Pedrós-Alió, C, Sebastián, M, Labadie, K, Paoli, L, Ibarbalz, FM, Zinger, L, Churcheward, B, Babin, M, Bork, P, Boss, E, Cochrane, G, De Vargas, C, Gorsky, G, Grimsley, N, Guidi, L, Hingamp, P, Iudicone, D, Jaillon, O, Kandels, S, Not, F, Ogata, H, Pesant, S, Poulton, N, Raes, J, Sardet, C, Speich, S, Stemmann, L, Sullivan, MB, Chaffron, S, Eveillard, D, Karsenti, E, Sunagawa, S, Wincker, P, Karp-Boss, L, Bowler, C, Acinas, SG. 2021. Compendium of 530 metagenome-assembled bacterial and archaeal genomes from the polar Arctic Ocean. Nature Microbiology 6(12): 1561–1574. DOI: 10.1038/s41564-021-00979-9.

Salcher, MM, Neuenschwander, SM, Posch, T, Pernthaler, J. 2015. The ecology of pelagic freshwater methylotrophs assessed by a high-resolution monitoring and isolation campaign. The ISME Journal 9(11): 2442–2453. DOI: 10.1038/ismej.2015.55.

Sayers, EW, Bolton, EE, Brister, JR, Canese, K, Chan, J, Comeau, DC, Connor, R, Funk, K, Kelly, C, Kim, S, Madej, T, Marchler-Bauer, A, Lanczycki, C, Lathrop, S, Lu, Z, Thibaud-Nissen, F, Murphy, T, Phan, L, Skripchenko, Y, Tse, T, Wang, J, Williams, R, Trawick, BW, Pruitt, KD, Sherry, ST. 2022. Database resources of the National Center for Biotechnology Information. Nucleic Acids Research 50(D1): D20–D26. DOI: 10.1093/nar/gkab1112.

Schembri, S, Bouchard, C, Pontbriand, T, Maps, F. 2023. Seasonality of zooplankton communities in the Hudson Bay system. Elementa: Science of the Anthropocene 11(1): 00113. DOI: 10.1525/elementa.2022.00113.

Schiffrine, N, Dhifallah, F, Dionne, K, Poulin, M, Lessard, S, Rochon, A, Gosselin, M. 2024. Microbial plankton occurrence database in the North American Arctic region: Synthesis of recent diversity of potentially toxic and/or harmful algae. Earth System Science Data 16(12): 5681–5701. DOI: 10.5194/essd-16-5681-2024.

Schloss, PD. 2024. Rarefaction is currently the best approach to control for uneven sequencing effort in amplicon sequence analyses. mSphere 9(2): e00354–23. DOI: 10.1128/msphere.00354-23.

Sipler, RE, Kellogg, CTE, Connelly, TL, Roberts, QN, Yager, PL, Bronk, DA. 2017. Microbial community response to terrestrially derived dissolved organic matter in the coastal Arctic. Frontiers in Microbiology 8: 1018. DOI: 10.3389/fmicb.2017.01018.

Six, C, Thomas, JC, Garczarek, L, Ostrowski, M, Dufresne, A, Blot, N, Scanlan, DJ, Partensky, F. 2007. Diversity and evolution of phycobilisomes in marine *Synechococcus* spp.: A comparative genomics study. Genome Biology 8(12): R259. DOI: 10.1186/gb-2007-8-12-r259.

Stadler, M, del Giorgio, PA. 2022. Terrestrial connectivity, upstream aquatic history and seasonality shape bacterial community assembly within a large boreal aquatic network. The ISME Journal 16(4): 937–947. DOI: 10.1038/s41396-021-01146-y.

Stoecker, DK, Lavrentyev, PJ. 2018. Mixotrophic plankton in the polar seas: A pan-Arctic review. Frontiers in Marine Science 5: 292. DOI: 10.3389/fmars.2018.00292.

Straneo, F, Saucier, F. 2008. The outflow from Hudson Strait and its contribution to the Labrador Current. Deep-Sea Research Part I 55(8): 926–946. DOI: 10.1016/j.dsr.2008.03.012.

Strunecký, O, Ivanova, AP, Mareš, J. 2023. An updated classification of cyanobacterial orders and families based on phylogenomic and polyphasic analysis. Journal of Phycology 59(1): 12–51. DOI: 10.1111/jpy.13304.

Teeling, H, Fuchs, BM, Becher, D, Klockow, C, Gardebrecht, A, Bennke, CM, Kassabgy, M, Huang, S, Mann, AJ, Waldmann, J, Weber, M, Klindworth, A, Otto, A, Lange, J, Bernhardt, J, Reinsch, C, Hecker, M, Peplies, J, Bockelmann, FD, Callies, U, Gerdts, G, Wichels, A, Wiltshire, KH, Glöckner, FO, Schweder, T, Amann, R. 2012. Substrate-controlled succession of marine bacterioplankton populations induced by a phytoplankton bloom. Science 336(6081): 608–611. DOI: 10.1126/science.1218344.

Teeling, H, Fuchs, BM, Bennke, CM, Krüger, K, Chafee, M, Kappelmann, L, Reintjes, G, Waldmann, J, Quast, C, Glöckner, FO, Lucas, J, Wichels, A, Gerdts, G, Wiltshire, KH, Amann, RI. 2016. Recurring patterns in bacterioplankton dynamics during coastal spring algae blooms. eLife 5: e11888. DOI: 10.7554/eLife.11888.

Thiele, S, Vader, A, Thomson, S, Saubrekka, K, Petelenz-Kurdziel, E, Armo, HR, Müller, O, Olsen, L, Bratbak, G, Øvreås, L. 2023. The summer bacterial and archaeal community composition of the northern Barents Sea. Progress in Oceanography 215: 103054. DOI: 10.1016/j.pocean.2023.103054.

Tisserand, L, Dadaglio, L, Intertaglia, L, Catala, P, Panagiotopoulos, C, Obernosterer, I, Joux, F. 2020. Use of organic exudates from two polar diatoms by bacterial isolates from the Arctic Ocean. Philosophical Transactions of the Royal Society A 378(2181): 20190356. DOI: 10.1098/rsta.2019.0356.

Unfried, F, Becker, S, Robb, CS, Hehemann, JH, Markert, S, Heiden, SE, Hinzke, T, Becher, D, Reintjes, G, Krüger, K, Avcı, B, Kappelmann, L, Hahnke, RL, Fischer, T, Harder, J, Teeling, H, Fuchs, B, Barbeyron, T, Amann, RI, Schweder, T. 2018. Adaptive mechanisms that provide competitive advantages to marine Bacteroidetes during microalgal blooms. The ISME Journal 12(12): 2894–2906. DOI: 10.1038/s41396-018-0243-5.

von Scheibner, M, Herlemann, DPR, Lewandowska, AM, Jürgens, K. 2018. Phyto- and bacterioplankton during early spring conditions in the Baltic Sea and response to short-term experimental warming. Frontiers in Marine Science 5: 231. DOI: 10.3389/fmars.2018.00231.

Yurkowski, DJ, Johnson, KF, Carvalho, PC, Johnson, MW, Prodaehl, AW, Mackey, T, Dawson, K, Candlish, L, Capelle, DW, Dunmall, K, Harris, LN, Ferguson, SH, Marcoux, M, McNicholl, D, Mosnier, A, Niemi, A, O’Brien, J, Pierrejean, M, Watt, C. 2023. Biophysical and ecological overview summary of the Qikiqtait study area and adjacent waters. Ottawa: Fisheries and Oceans Canada. Canadian Technical Report of Fisheries and Aquatic Sciences 3565. Available at https://publications.gc.ca/site/eng/9.927290/publication.html.

