## Supplementary Material for "Microbial plankton in southern Hudson Bay and James Bay in late summer"

**Contents:** Figure S1; Figure S2; Tables S1–S4.

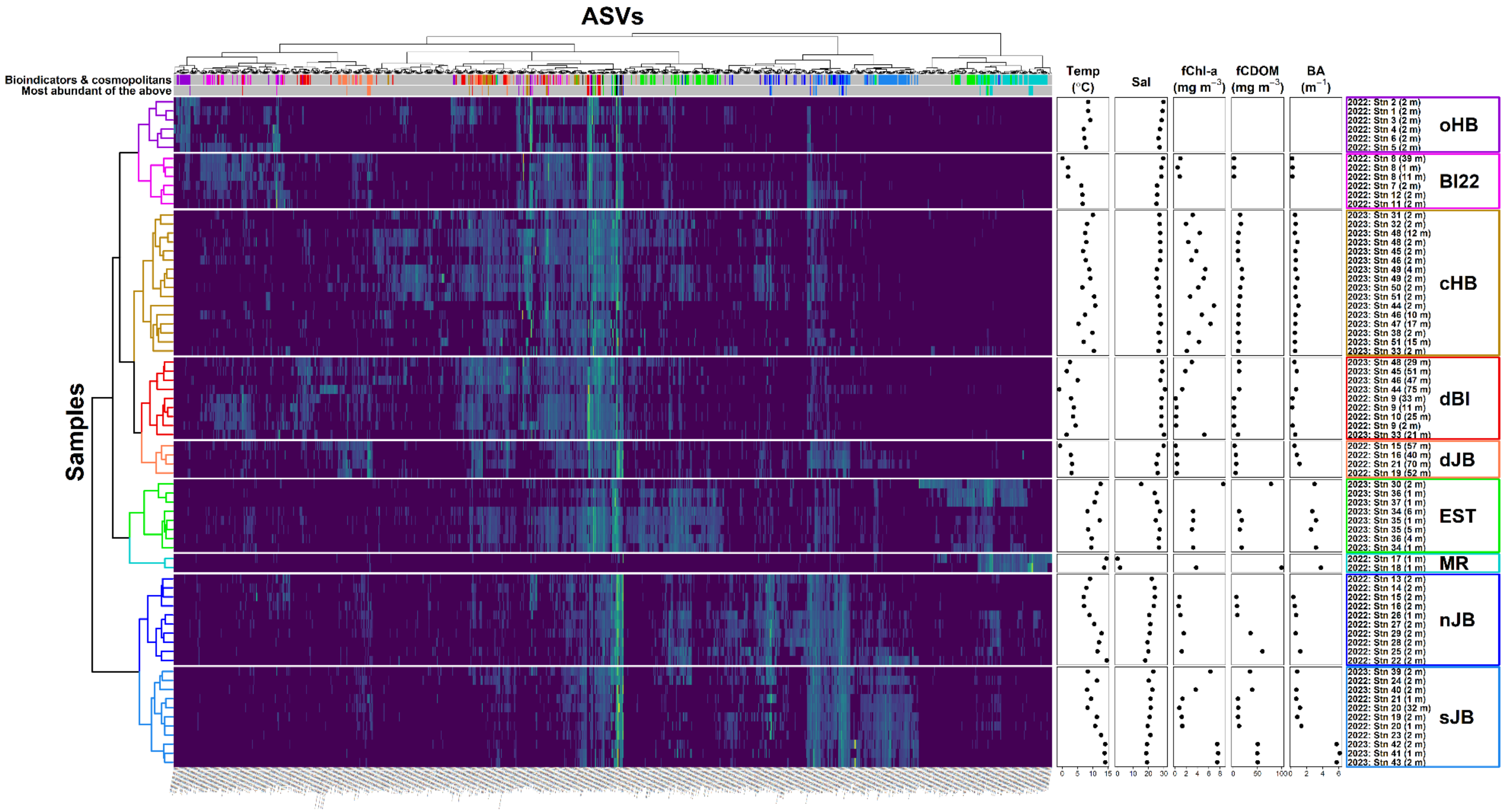


**Figure S1. Heatmap of transformed relative abundance of non-rare eukaryotic and prokaryotic ASVs across samples collected in SHJB.** Samples, expressed with year, station, and depth of sampling, are indicated on the right. Environmental parameters associated with the samples – temperature (Temp), salinity (Sal), chlorophyll-a fluorescence (fChl-a), CDOM fluorescence (fCDOM), and beam attenuation (BA) – are also presented on the right. The dendrogram on the left represents hierarchical clustering of samples. The panels above the heatmap highlight bioindicators (color designates the cluster each bioindicator has an affinity for) and cosmopolitans (black). The top panel highlights all bioindicator and cosmopolitan ASVs, and the bottom panel highlights only the ASVs reaching $\geq 1.5\%$ mean relative abundance in their indicator cluster, or across all samples if cosmopolitan.

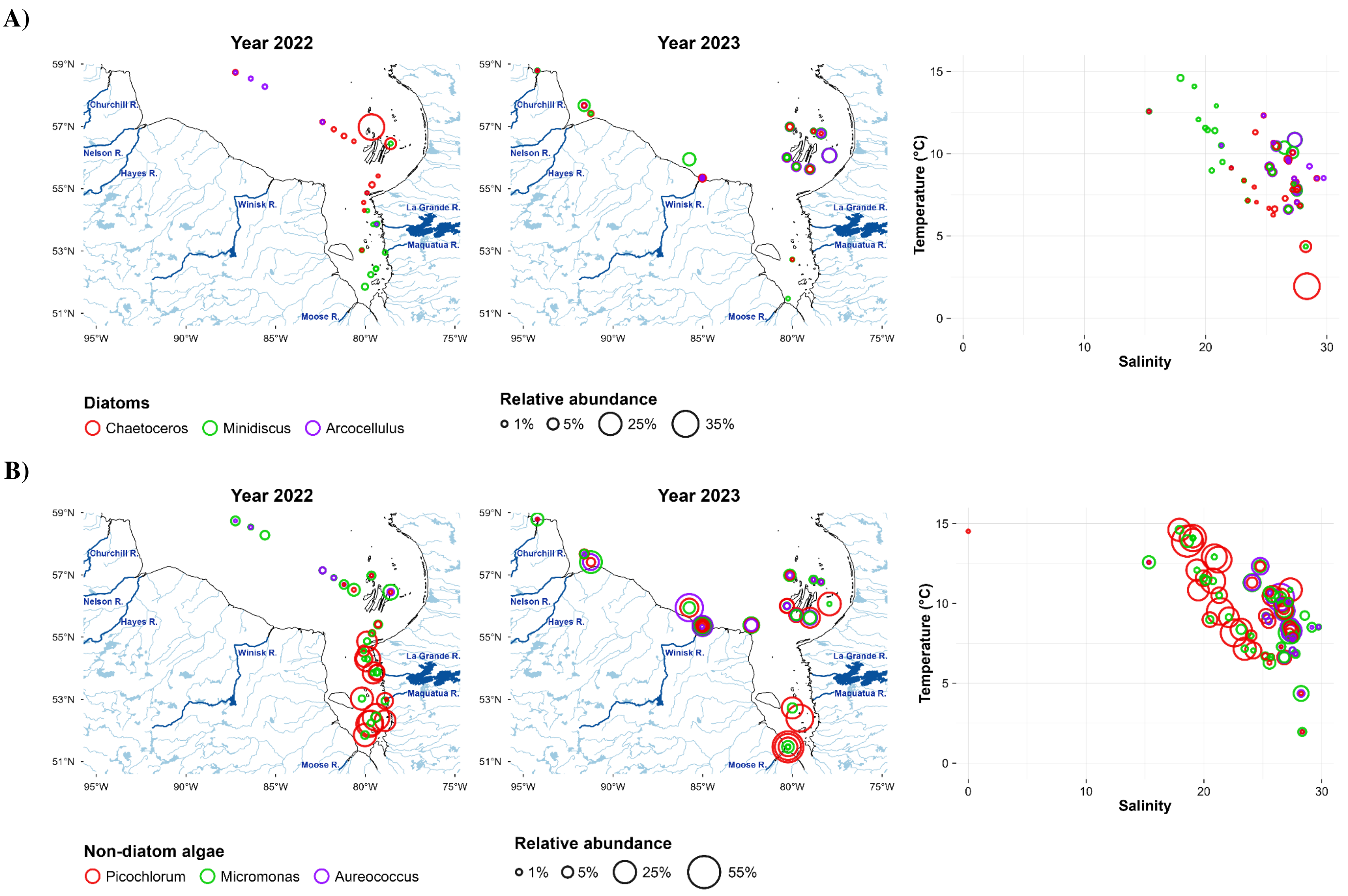


**Figure S2. Primary producers in surface waters (1-6 m) of SHJB in August: A) diatoms; B) non-diatom algae; C) non-parasitic dinoflagellates; D) cyanobacteria.** Relative abundances are shown for 2022 and 2023, and in relation to temperature and salinity.

C)

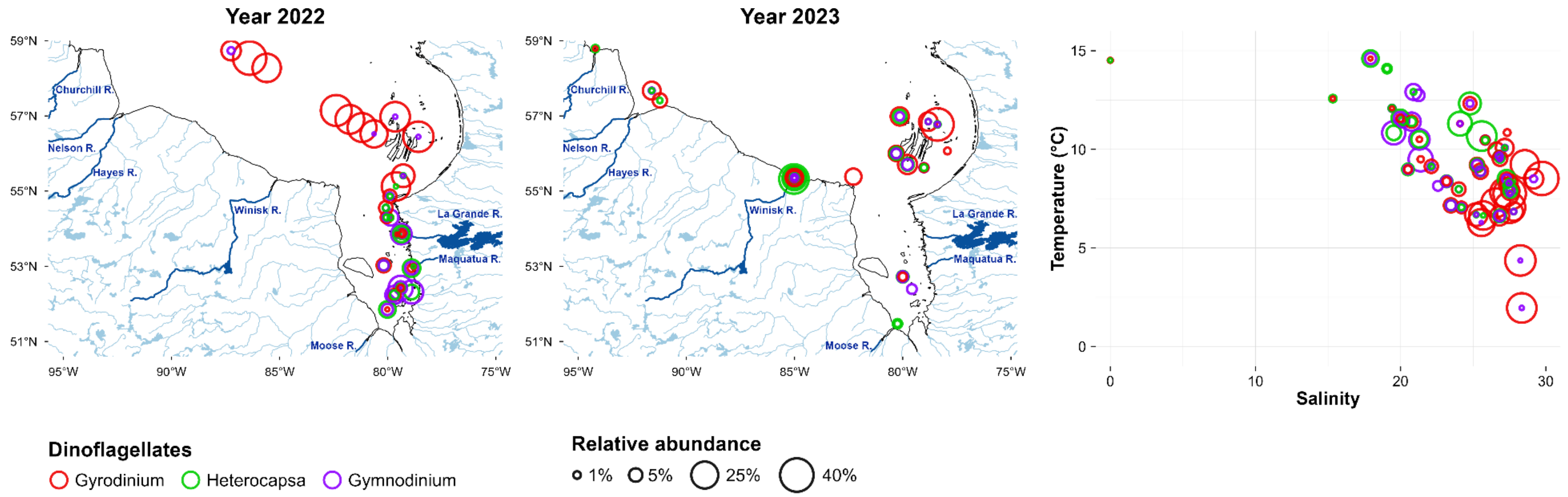


D)

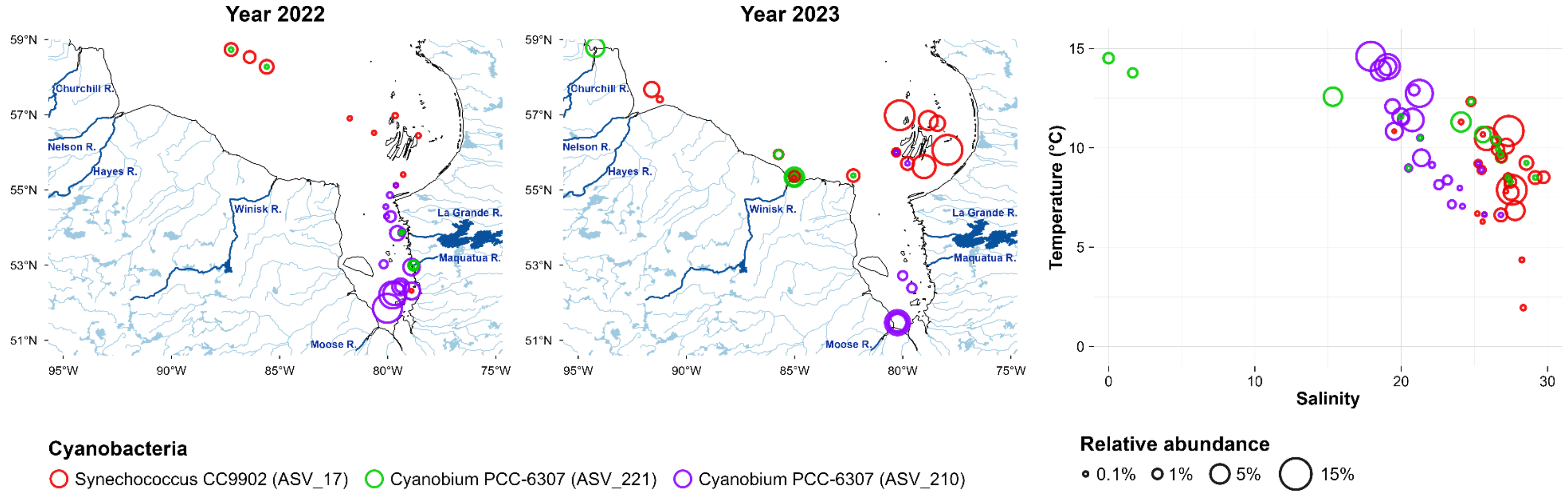


**Figure S2.** (continued) **C)** non-parasitic dinoflagellates; **D)** cyanobacteria.

**Table S1. Environmental properties of sample clusters in SHJB.** Values are medians [ranges] per cluster. Asterisk signifies values, where some samples in the cluster (no more than half) had no measurements.

| Cluster | Year | Depth (m) | Distance to coast (km) | Temperature (°C) | Salinity | fChl-a (mg m$^{-3}$) | fCDOM (mg m$^{-3}$) | Beam Attenuation (m$^{-1}$) |
|---|---|---|---|---|---|---|---|---|
| oHB (offshore Hudson Bay) | 2022 | 2.0 | 187.0 [75.0-243.0] | 8.2 [7.1-9.2] | 28.0 [26.5-29.7] | NA | NA | NA |
| BI22 (the Belcher Islands) | 2022 | 2.0 [1.4-38.6] | 18.0 [8.0-43.0] | 4.1 [0.1-6.7] | 27.0 [25.2-29.5] | 0.9* [0.5-1.0] | 1.6* [1.6-1.6] | 0.2* [0.2-0.3] |
| cHB (coastal Hudson Bay) | 2023 | 2.0 [1.5-16.6] | 20.0 [9.0-53.0] | 8.0 [5.3-10.9] | 27.3 [25.3-28.1] | 4.0 [2.0-6.9] | 11.3 [9.7-18.5] | 0.6 [0.5-1.0] |
| dBI (the Belcher Islands, subsurface) | 2022-2023 | 29.0 [1.9-75.0] | 17.0 [1.0-32.0] | 2.9 [-1.0-5.1] | 28.5 [27.8-30.7] | 0.8* [0.1-5.2] | 5.9* [1.6-12.4] | 0.5* [0.2-0.8] |
| dJB (James Bay, subsurface) | 2022 | 54.3 [40.1-70.1] | 18.0 [13.0-36.0] | 2.9 [-0.7-3.3] | 26.1 [25.3-29.8] | 0.3 [0.2-0.4] | 5.3 [2.7-6.2] | 0.8* [0.5-1.1] |
| EST (the Churchill and Winisk estuaries) | 2023 | 1.7 [1.0-6.0] | 6.0 [0.0-7.0] | 10.2 [8.3-12.6] | 26.2 [15.3-27.5] | 3.2* [3.0-8.6] | 17.5* [12.2-78.2] | 3.0* [2.6-3.2] |
| MR (the Maquatua River) | 2022 | 1.0 | 1.5 [1.0-2.0] | 14.1 [13.8-14.5] | 0.8 [0.0-1.6] | 3.8* | 99.7* | 3.8* |
| nJB (northern James Bay) | 2022 | 2.0 [1.4-2.2] | 20.0 [2.0-36.0] | 9.8 [7.1-14.6] | 21.1 [17.9-24.2] | 1.0* [0.7-1.6] | 8.5* [6.8-60.2] | 0.6* [0.3-1.2] |
| sJB (southern James Bay) | 2022-2023 | 2.0 [1.2-31.8] | 13.0 [12.0-31.0] | 11.4 [8.1-14.1] | 20.8 [18.6-23.2] | 3.7* [0.8-7.7] | 34.5* [9.8-50.7] | 1.2* [0.7-6.1] |
| Kruskal-Wallis p | NA | $9.8 \times 10^{-5}$ | $9.8 \times 10^{-6}$ | $3.0 \times 10^{-8}$ | $6.5 \times 10^{-9}$ | $3.5 \times 10^{-4}$ | $5.5 \times 10^{-4}$ | $5.5 \times 10^{-5}$ |

**Table S2. Significant post-hoc Dunn's pairwise comparisons of environmental parameters among SHJB sample clusters.** Only comparisons with Benjamini-Hochberg-adjusted $p < 0.05$ are shown. The ">" symbol points towards the cluster with the lower median.

| Parameter | Direction of difference | BH-adjusted p-value |
|---|---|---|
| Beam Attenuation ($m^{-1}$) | sJB > BI22 | $2.7 \times 10^{-3}$ |
| Beam Attenuation ($m^{-1}$) | EST > BI22 | $3.1 \times 10^{-3}$ |
| Beam Attenuation ($m^{-1}$) | EST > dBI | $4.1 \times 10^{-3}$ |
| Beam Attenuation ($m^{-1}$) | sJB > dBI | $4.3 \times 10^{-3}$ |
| Beam Attenuation ($m^{-1)}$ | EST > cHB | $1.5 \times 10^{-2}$ |
| Beam Attenuation ($m^{-1}$) | sJB > cHB | $1.8 \times 10^{-2}$ |
| Beam Attenuation ($m^{-1}$) | MR > BI22 | $3.5 \times 10^{-2}$ |
| Beam Attenuation ($m^{-1}$) | EST > nJB | $4.9 \times 10^{-2}$ |
| Depth (m) | dJB > MR | $7.8 \times 10^{-3}$ |
| Depth (m) | dJB > sJB | $1.8 \times 10^{-2}$ |
| Depth (m) | dJB > EST | $2.1 \times 10^{-2}$ |
| Depth (m) | dJB > cHB | $2.2 \times 10^{-2}$ |
| Depth (m) | dBI > MR | $2.4 \times 10^{-2}$ |
| Distance to coast (km) | BI22 > sJB | $9.1 \times 10^{-4}$ |
| Distance to coast (km) | sJB > EST | $2.6 \times 10^{-3}$ |
| Salinity | dBI > sJB | $2.6 \times 10^{-6}$ |
| Salinity | dBI > nJB | $9.4 \times 10^{-6}$ |
| Salinity | oHB > sJB | $8.3 \times 10^{-4}$ |
| Salinity | cHB > sJB | $9.2 \times 10^{-4}$ |
| Salinity | dBI > MR | $1.1 \times 10^{-3}$ |
| Salinity | oHB > nJB | $1.4 \times 10^{-3}$ |
| Salinity | cHB > nJB | $2.5 \times 10^{-3}$ |
| Salinity | BI22 > sJB | $6.7 \times 10^{-3}$ |
| Salinity | oHB > MR | $6.7 \times 10^{-3}$ |
| Salinity | dBI > EST | $8.5 \times 10^{-3}$ |
| Salinity | BI22 > nJB | $1.2 \times 10^{-2}$ |
| Salinity | cHB > MR | $1.8 \times 10^{-2}$ |
| Salinity | BI22 > MR | $2.3 \times 10^{-2}$ |
| Salinity | dJB > sJB | $4.0 \times 10^{-2}$ |
| fCDOM (mg $m^{-3}$) | nJB > BI22 | $1.2 \times 10^{-2}$ |
| fCDOM (mg $m^{-3}$) | dJB > BI22 | $2.5 \times 10^{-2}$ |
| fCDOM (mg $m^{-3}$) | sJB > BI22 | $2.7 \times 10^{-2}$ |
| fCDOM (mg $m^{-3}$) | MR > BI22 | $2.7 \times 10^{-2}$ |
| fCDOM (mg $m^{-3}$) | nJB > dBI | $2.9 \times 10^{-2}$ |
| fChl-a (mg $m^{-3}$) | cHB > dJB | $1.3 \times 10^{-2}$ |
| fChl-a (mg $m^{-3}$) | cHB > dBI | $1.9 \times 10^{-2}$ |
| fChl-a (mg $m^{-3}$) | sJB > dJB | $2.5 \times 10^{-2}$ |
| fChl-a (mg $m^{-3}$) | EST > dJB | $3.4 \times 10^{-2}$ |

**Table S3. Significant post-hoc Dunn's pairwise comparisons of Shannon diversity indices among SHJB sample clusters.** Only comparisons with Benjamini-Hochberg-adjusted $p < 0.05$ are shown. The ">" symbol points towards the cluster with the lower median.

| Domain | Direction of difference | BH-adjusted p-value |
|---|---|---|
| Eukaryota | EST > sJB | $1.3 \times 10^{-3}$ |
| Eukaryota | dBI > sJB | $1.3 \times 10^{-2}$ |
| Eukaryota | cHB > sJB | $1.8 \times 10^{-2}$ |
| Eukaryota | EST > BI22 | $2.0 \times 10^{-2}$ |
| Prokaryota | EST > cHB | $3.5 \times 10^{-3}$ |
| Prokaryota | EST > oHB | $4.7 \times 10^{-3}$ |
| Prokaryota | EST > BI22 | $5.1 \times 10^{-3}$ |
| Prokaryota | dJB > oHB | $1.2 \times 10^{-2}$ |
| Prokaryota | sJB > oHB | $1.3 \times 10^{-2}$ |
| Prokaryota | sJB > cHB | $1.5 \times 10^{-2}$ |
| Prokaryota | dJB > BI22 | $1.6 \times 10^{-2}$ |
| Prokaryota | dJB > cHB | $1.7 \times 10^{-2}$ |
| Prokaryota | sJB > BI22 | $1.9 \times 10^{-2}$ |
| Prokaryota | MR > oHB | $2.1 \times 10^{-2}$ |
| Prokaryota | dBI > oHB | $4.4 \times 10^{-2}$ |
| Prokaryota | MR > cHB | $4.5 \times 10^{-2}$ |
| Prokaryota | nJB > oHB | $4.6 \times 10^{-2}$ |
| Prokaryota | MR > BI22 | $4.7 \times 10^{-2}$ |

**Table S4. Significant Spearman correlations between Shannon diversity indices and spatiotemporal or environmental parameters in SHJB.** Only correlations with $p < 0.001$ are shown. n is the number of samples, and $\rho$ is Spearman's rank correlation coefficient.

| Domain | Parameter | n | $\rho$ | p-value |
|---|---|---|---|---|
| Prokaryota | Distance to coast [km] | 67 | -0.46915005 | $6.2 \times 10^{-5}$ |
| Prokaryota | Latitude [deg] | 67 | -0.43626967 | $2.2 \times 10^{-4}$ |
| Prokaryota | Bot. Depth [m] | 67 | -0.41967746 | $4.1 \times 10^{-4}$ |
